# Maturing Mycobacterial Peptidoglycan Requires Non-canonical Crosslinks to Maintain Shape

**DOI:** 10.1101/291823

**Authors:** Catherine Baranowski, Michael A. Welsh, Lok-To Sham, Haig A. Eskandarian, Hoong C. Lim, Karen J. Kieser, Jeffrey C. Wagner, John D. McKinney, Georg E. Fantner, Thomas R. Ioerger, Suzanne Walker, Thomas G. Bernhardt, Eric J. Rubin, E. Hesper Rego

**Affiliations:** Department of Immunology and Infectious Disease, Harvard T. H. Chan School of Public Health, Boston, MA 02115, USA; Department of Microbiology and Immunobiology, Harvard Medical School, Boston, MA 02115, USA; Department of Microbiology and Immunology, National University of Singapore, 117545 Singapore; School of Life Sciences, Swiss Federal Institute of Technology in Lausanne (EPFL), 1015 Lausanne, Switzerland; School of Engineering, Swiss Federal Institute of Technology in Lausanne (EPFL), 1015 Lausanne, Switzerland; Department of Computer Science and Engineering, Texas A & M University, College Station, Texas, USA; Department of Microbial Pathogenesis, Yale University School of Medicine, New Haven, Connecticut, 06510, USA

## Abstract

In most well studied rod-shaped bacteria, peptidoglycan is primarily crosslinked by penicillin binding proteins (PBPs). However, in mycobacteria, L,D-transpeptidase (LDT)-mediated crosslinks are highly abundant. To elucidate the role of these unusual crosslinks, we characterized mycobacterial cells lacking all LDTs. We find that LDT-mediated crosslinks are required for rod shape maintenance specifically at sites of aging cell wall, a byproduct of polar elongation. Asymmetric polar growth leads to a non-uniform distribution of these two types of crosslinks in a single cell. Consequently, in the absence of LDT-mediated crosslinks, PBP-catalyzed crosslinks become more important. Because of this, *Mycobacterium tuberculosis* (Mtb) is more rapidly killed using a combination of drugs capable of PBP- and LDT-inhibition. Thus, knowledge about the single-cell distribution of drug targets can be exploited to more effectively treat this pathogen.

## Introduction

Peptidoglycan (PG) is an essential component of all bacterial cells (Vollmer, Blanot, & de Pedro, 2008a), and the target of many antibiotics. PG consists of linear glycan strands crosslinked by short peptides to form a continuous molecular cage surrounding the plasma membrane. This structure maintains cell shape and protects the plasma membrane from rupture. Our understanding of PG is largely derived from studies on laterally growing model rod-shaped bacteria like *Escherichia coli* and *Bacillus subtilis* (*Figure 1 - figure supplement 1A*). In these organisms, new PG is constructed along the lateral side wall by the concerted effort of glycoslytransferases, which connect the glycan of a new PG subunit to the existing mesh, and transpeptidases, which link peptide side chains. An actin-like protein, MreB, positions this multi-protein complex along the short axis of the cell so that glycan strands are inserted circumferentially, creating discontinuous hoops of PG around the cell (Domínguez-Escobar et al., 2011; Garner et al., 2011). This orientation of PG creates a mechanical anisotropy that is responsible for rod shape (Hussain et al., 2018).

**Figure 1:**
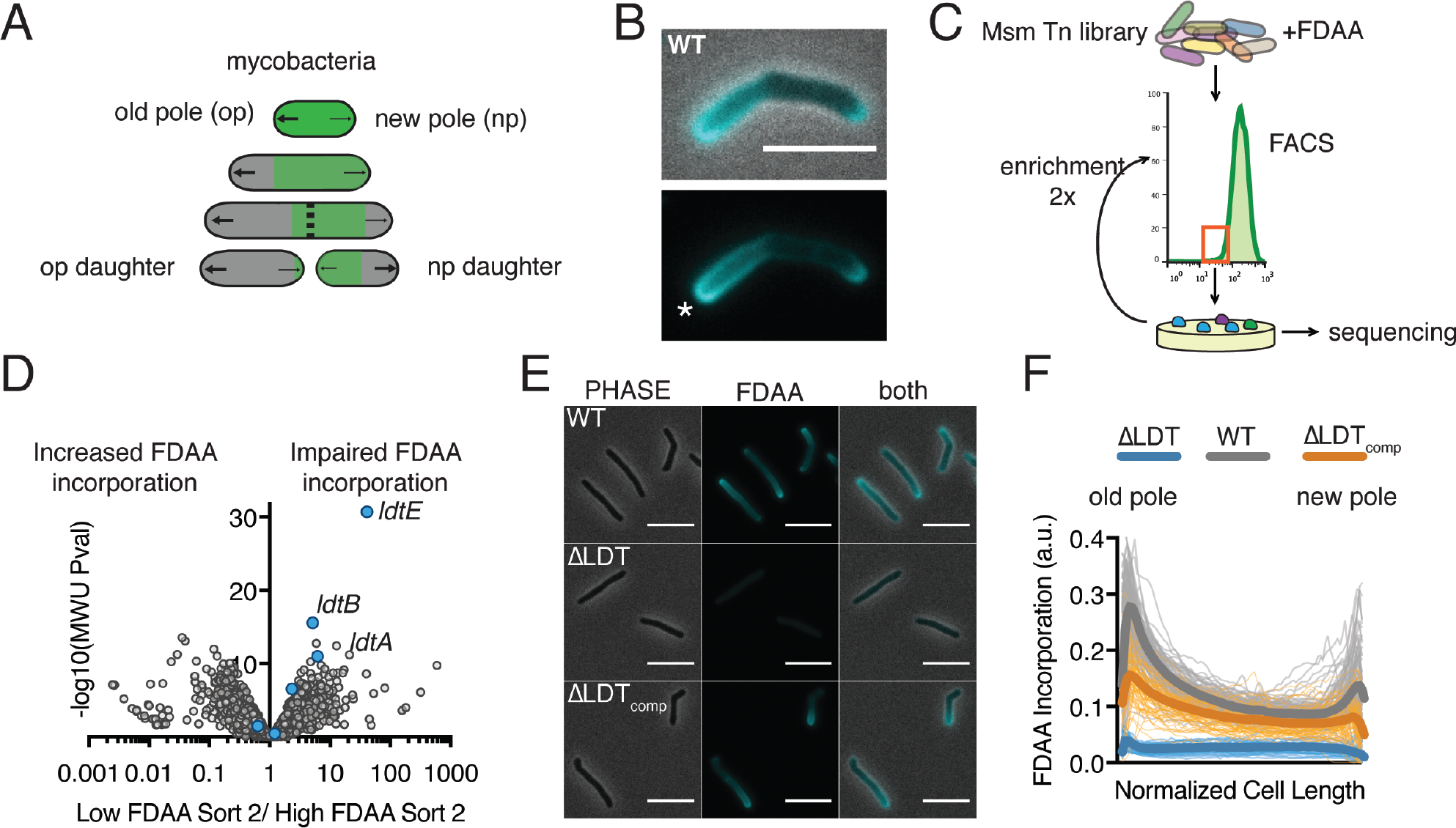
FDAAs are incorporated asymmetrically by LD-transpeptidases. **(A)** Schematic of mycobacterial asymmetric polar growth. Green, old cell wall; grey, new material; dotted line, septum; large arrows, old pole growth; small arrows, new pole growth. **(B)** FDAA incorporation in log-phase WT Msm cell after 2-minute incubation. Scale bar=5µm. Old pole marked with (*). **(C)** Schematic of Fluorescence Activated Cell Sorting (FACS)-based FDAA transposon library enrichment. An Msm transposon (Tn) library was stained with FDAAs, the dimmest and brightest cells were sorted, grown, sorted again to enrich for transposon mutants that are unable or enhanced for FDAA incorporation. **(D)** Results from 1C screen. For each gene, the contribution to low or high staining population was calculated from transposon reads per gene. Plotted is the ratio of the population contribution from the second sort of low FDAA staining (L2) over the second sort of high FDAA staining (H2) cells compared to the Mann-Whitney *U* P-value. **(E)** Representative image of FDAA incorporation in log-phase WT, ∆LDT and ∆LDT_comp_ cells. Scale bar= 5μm. **(F)** Profiles of FDAA incorporation in log-phase WT (N=98), ∆LDT (N=40), and ∆LDT_comp_ (N=77) cells. Thick lines represent mean incorporation profile, thin lines are FDAA incorporation in single cells.

However, not all rod-shaped bacteria encode MreB. In fact, there are important differences between model bacteria and Actinobacteria like mycobacteria, a genus of rod-shaped bacteria that includes the major human pathogen *Mycobacterium tuberculosis* (Mtb). In mycobacteria, new PG is inserted at the cell poles (at unequal amounts based on pole age), rather than along the lateral walls (*Figure 1A*). Additionally, mycobacteria are missing several factors, including MreB, that are important for cell elongation (Kieser & Rubin, 2014). Furthermore, in *E. coli* and *B. subtilis* the vast majority (>90%) of the peptide linkages are created by D,D-transpeptidases known as penicillin binding proteins (PBPs) (Pisabarro, de Pedro, & Vázquez, 1985). PBPs, the targets of most b-lactams, link the 4^th^ amino acid of one peptide side chain to the 3^rd^ amino acid of another, forming 4-3 crosslinks (*Figure 1-figure supplement 1B*). Peptide linkages can also be catalyzed by L,D-transpeptidases (LDTs), which link peptide side chains by the 3^rd^ amino acid forming 3-3 linkages. In mycobacteria, these 3-3 crosslinks, are highly abundant, accounting for ~60% of linkages (Kumar et al., 2012) (*Figure 1-figure supplement 1B*). Because PG has been most well studied in bacteria where 3-3 crosslinks are rare, the role of these linkages, and the enzymes that catalyze their formation, is poorly understood.

Tuberculosis remains an enormous global health problem, in part, because treating even drug susceptible disease is difficult. The standard regimen includes a cocktail of four drugs given over six months. Treatment of drug-resistant Mtb is substantially longer and includes combinations of up to seven drugs (“Global Tuberculosis Report 2017,” 2017). While some of the most important anti-mycobacterials target cell wall synthesis, surprisingly, drugs that target PG are not part of the core treatment for either drug-susceptible or drug-resistant disease. However, carbapenems, b-lactam antibiotics that potently inhibit LDTs *in vitro*, are effective against drug resistant Mtb *in vitro* and drug sensitive Mtb in patients (Diacon et al., 2016; Hugonnet, Tremblay, Boshoff, Barry, & Blanchard, 2009).

But, why are LDTs important in mycobacteria? To explore this, we constructed strains that lack the ability to form 3-3 crosslinks. We find that 3-3 crosslinks are formed in maturing peptidoglycan and that they are necessary to stabilize the cell wall and prevent lysis. Cells that lose the ability to synthesize 3-3 crosslinks have increased dependence on 4-3 crosslinking. Thus, simultaneous inhibition of both processes results in rapid cell death.

## Results

### Fluorescent D-amino acids are incorporated asymmetrically by LD-transpeptidases

PG uniquely contains D-amino acids, which can be labeled with fluorescent probes (fluorescent D-amino acids, FDAAs) to visualize PG synthesis in live bacterial cells (Kuru et al., 2012). When we incubated *M. smegmatis* (Msm), a non-pathogenic relative of Mtb, with FDAAs for a short 2-minute pulse (< 2% of Msm’s generation time) we observed incorporation at both poles, the sites of new PG insertion in mycobacteria (*Figure 1A, B*) (Aldridge et al., 2012). However, we also saw a gradient of fluorescence along the sidewalls, extending from the old pole (the previously established growth pole) that fades to a minimum at roughly mid-cell as it reaches the new pole (the pole formed at the last cell division) (*Figure 1B*).

To identify the enzymes responsible for this unexpected pattern of lateral cell wall FDAA incorporation, we performed a fluorescence-activated cell sorting (FACS)-based transposon screen (*Figure 1C*). Briefly, we stained an Msm transposon library with FDAA and repeatedly sorted the least fluorescent 12.5% of the population by FACS. After each sort we regrew cells, extracted gDNA and used deep sequencing to map the location of the transposons found in the low staining population.

From this screen, we identified three LDTs (*ldtA - MSMEG_3528, ldtB - MSMEG_4745, ldtE* - *MSMEG_0233*) (*Figure 1D*) that were primarily responsible for FDAA incorporation. Deleting these three LDTs significantly reduced FDAA incorporation (*Figure 1-figure supplement 2A, B*). To further investigate the physiological role of LDTs, we constructed a strain lacking all 6 LDTs (D*ldtAEBCGF*, hereafter DLDT). FDAA incorporation and 3-3 crosslinking are both nearly abolished in DLDT cells and can be partially restored by complementation with a single LDT (*ldtE*-mRFP; DLDT_comp_) (*Figure 1E, F, Figure 1-figure supplement 2C, 3*). Thus, as might be the case in *Bdellovibrio* (Kuru et al., 2017), FDAA incorporation in Msm is primarily LDT-dependent. LDTs have previously been shown to exchange non-canonical D-amino acids onto PG tetrapeptides in *Vibrio cholera* (Cava, de Pedro, Lam, Davis, & Waldor, 2011).

### LDT catalyzed 3-3 crosslinks are required for rod shape maintenance at aging cell wall

As deletion of a subset of LDTs in Msm produces morphologic changes (Sanders, Wright, & Pavelka, 2014), we visualized ΔLDT cells by time-lapse microscopy. We observed that a subpopulation of cells loses rod shape progressively over time, resulting in localized spherical blebs (*Figure 2A - top, Figure 2 - figure supplement 1A, Figure 2 - video 1*). Complemented cells are able to maintain rod shape (*Figure 2 - figure supplement 1B*). We reasoned that localized loss of rod shape may occur for two reasons:1) spatially-specific loss of cell wall integrity and/or 2) cell wall deformation due to uncontrolled, local PG synthesis. If the first hypothesis were true, high osmolarity should protect cells against forming blebs. Indeed, switching cells from iso- to high-osmolarity prevented bleb formation over time (*Figure 2A, bottom, Figure 2-video 2*). To test the second hypothesis, we stained ΔLDT cells with an amine-reactive dye, and observed outgrowth of new, unstained material (*Figure 2B*). Blebs retained stain, indicating a lack of new cell wall synthesis in the region. Collectively, these results indicate that 3-3 crosslinks are required to counteract turgor pressure and maintain the rod shape of mycobacteria. This led us to hypothesize that bleb formation is a result of a local defect in cell wall rigidity.

**Figure 2:**
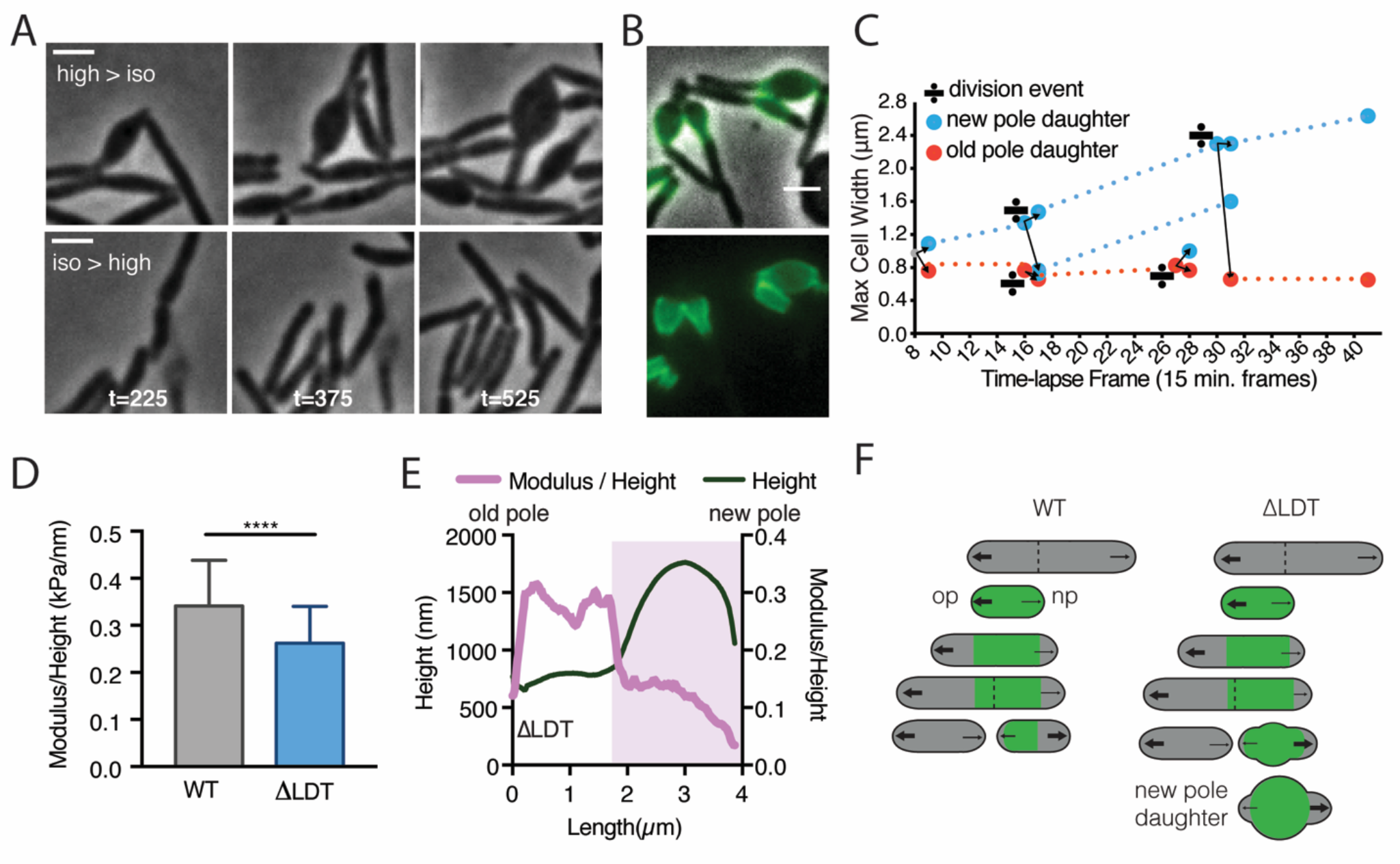
LDT catalyzed 3-3 crosslinks are required for rod shape maintenance at aging cell wall. **(A)** Msm ∆LDT time-lapse microscopy of cells switched from high- to iso- (top) osmolar media, or iso- to high osmolar media (bottom). (high=7H9+150mM sorbitol; iso=7H9). t= time in minutes post osmolarity switch. **(B)** ∆LDT cells were stained with Alexa 488 NHS-ester (green) to mark existing cell wall, washed, and visualized after outgrowth (unstained material). A, B scale bar=2µm. **(C)** Maximum cell width of ∆LDT cell lineages over time. Width of new pole daughters = blue circle; width of old pole daughters = orange circle. Division signs denote a division event. At each division, there are two arrows from the dividing cell leading to the resulting new and old pole daughter cell widths (blue and orange respectively). **(D)** Mean stiffness of WT (N=73) and ∆LDT (N=47) Msm cells as measured by atomic force microscopy. Mann-Whitney U P-Value **** < 0.0001. (**E)** Representative profile of cell height and height-normalized stiffness (modulus/height) in a single ∆LDT cell. Pink shaded portion highlights location of a bleb. **(F)** Model of rod shape loss in old cell wall of ∆LDT cells compared to WT. Green portions of the cell represents old cell wall; grey portion represents new cell wall. The larger arrows indicate more growth from the old pole, while smaller arrows show less relative growth from the new pole. Dotted lines represent septa. op= old pole, np=new pole.

To directly measure cell wall rigidity, we used atomic force microscopy (AFM) on live ΔLDT and WT cells. We measured the rigidity of cells in relation to their height. Generally, WT cells are stiffer than DLDT cells (*Figure 2D*). Blebs in DLDT cells can be identified by a sharp increase in height (*Figure 2E, pink shaded*). Since circumferential stress of the rod measured by AFM is proportional to the radius of the cell, and inversely proportional to the thickness of the cell wall (an immeasurable quantity by AFM), we used cell height, a proxy for radius, to normalize the stiffness measurement. We found that stiffness drops in the area of blebs (*Figure 2E, pink shaded*).

Why does loss of rod shape occur locally and only in a subpopulation of cells? Mycobacterial polar growth and division results in daughter cells with phenotypic differences (Aldridge et al., 2012). For example, the oldest cell wall is specifically inherited by the new pole daughter (*Figure 2-supplement 2A*, (Aldridge et al., 2012)). We hypothesized that the loss of rod shape might occur in specific progeny generated by cell division. Indeed, the daughter which inherited the new pole from the previous round of division, and the oldest cell wall, consistently lost rod shape over time, while the old pole daughter maintained rod shape (*Figure 2C, Figure 2-supplement 2B*). In addition, blebs localized to the oldest cell wall (*Figure 2B*), as visualized by pulse-chase labeling of the cell wall. Thus, 3-3 crosslinking is likely occurring in the oldest cell wall, which is non-uniformly distributed along a single cell and in the population via asymmetric polar growth and division. Taken together, these data suggest that LDTs act locally to reinforce aging PG and to maintain rod shape in a subpopulation of Msm cells - specifically, new pole daughters (*Figure 2F*).

### Mycobacteria are hypersensitive to PBP inactivation in the absence of LDTs

Our observations lead to the following model (*Figure 6A*): PBP-catalyzed 4-3 crosslinks are formed at the poles where new PG is inserted and where pentapeptide substrates reside. These newly synthesized 4-3 crosslinks are then gradually cleaved (by D,D-endopeptidases) as PG ages and moves toward the middle of the cell, leaving tetrapeptide substrates for LDTs to create 3-3 crosslinks. This is consistent with the FDAA incorporation pattern, which reflects the abundance of tetrapeptide substrates available for LDT exchange. Specifically, there are more available tetrapeptides near the poles and fewer near mid-cell, the site of older PG (*Figure 1F*). In the absence of LDTs to catalyze 3-3 crosslinks, old cell wall loses integrity and turgor pressure causes bleb formation.

This model predicts that DLDT cells should be even more dependent on 4-3 crosslinking than wild type cells. To test this hypothesis, we used TnSeq (Long et al., 2015) to identify genes required for growth in cells lacking LDTs (*Figure 3A*). We found that mutants of two PBPs, *pbpA (MSMEG_0031c)* and *ponA2 (MSMEG_6201)*, were recovered at significantly lower frequencies in DLDT cells (*Figure 3B*). Likewise, using allele swapping (Kieser, Boutte, et al., 2015b) (*Figure 3C, Figure 3-figure supplement 1A*), a technique that tests the ability of various alleles to support viability, we found that the transpeptidase (TP) activity of PonA1, which is non-essential in WT cells (Kieser, Boutte, et al., 2015b), becomes essential in DLDT cells (*Figure 3D*). Thus, cells that lack 3-3 crosslinks are more dependent on 4-3 crosslinking enzymes.

**Figure 3:**
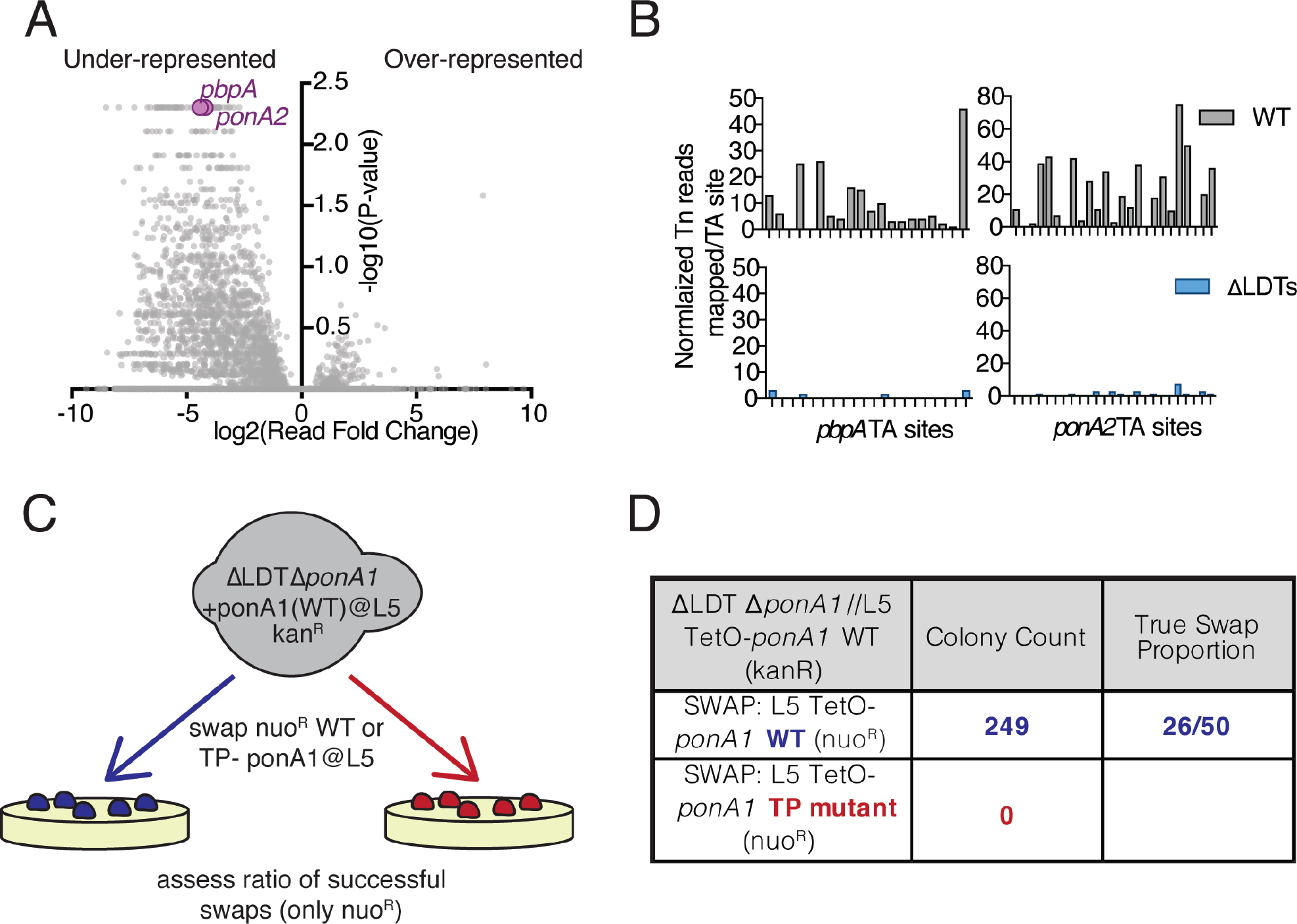
Mycobacteria are hypersensitive to PBP inactivation in the absence of LDTs. **(A)** Fold change in the number of reads for transposon insertion counts in ∆LDT cells compared to WT Msm. P-value is derived from a rank sum test (DeJesus, Ambadipudi, Baker, Sassetti, & Ioerger, 2015). **(B)** Transposon insertions per TA dinucleotide in *pbpA* and *ponA2* in WT (grey) and ∆LDT (blue) cells. **(C)** Schematic of L5 allele swapping experiment. Adapted from (Kieser, Boutte, et al., 2015b). **(D)** Results of WT or transpeptidase null *ponA1* allele swapping experiment in ∆LDT cells.

### Peptidoglycan synthesizing enzymes localize to differentially aged cell wall

Given our model, we hypothesized that enzymes catalyzing and processing different types of crosslinks should be differentially localized along the length of the cell. Specifically, we postulated that 4-3 crosslinking PBPs would localize at sites of new PG, while 4-3 cleaving D,D-endopeptidases and 3-3 crosslinking LDTs would localize to sites of older PG. To test whether 3-3 and 4-3 crosslinking enzymes localize differently, we visualized fluorescent fusions of a PBP (PonA1), and an LDT (LdtE), (*Figure 4A*). We found that PonA1-RFP largely localized to the old pole, where new PG is inserted (*Figure 4A, B, Figure 4 - video 1*). LdtE-mRFP localized farther from the poles, the sites of older PG (*Figure 4A, B, Figure 4 - video 2*). Thus, enzymes responsible for 4-3 and 3-3 crosslinks exhibit distinctive subcellular localizations, consistent with the model that they act on differentially aged PG.

**Figure 4:**
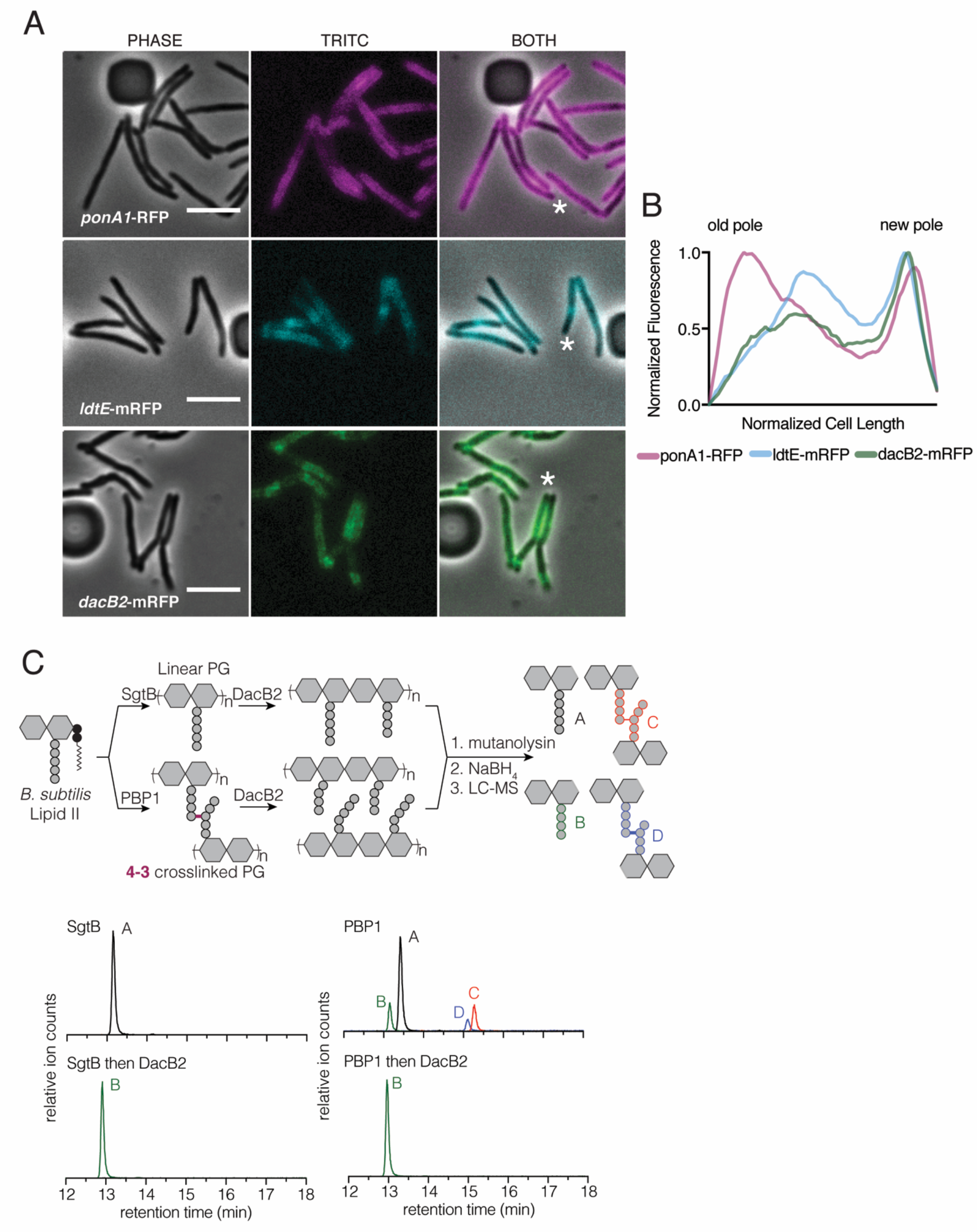
Peptidoglycan synthesizing enzymes localize to differentially aged cell wall. **(A)** Representative fluorescence images of PonA1-RFP (magenta), LdtE-mRFP (cyan), and DacB2-mRFP (green). Scale bars=5µm. **(B)** Average PonA1-RFP (N=24), LdtE-mRFP (N=23) or DacB2-mRFP (N=23) distribution in cells before division. **(C)** Schematic of the *in vitro* experiment to test D,D-carboxy- and D,D-endopeptidase activity of DacB2. Lipid II extracted from *B. subtilis* is first polymerized into linear (using SgtB) or crosslinked (using *B. subtilis* PBP1) peptidoglycan and then reacted with DacB2. The reaction products are analyzed by LC-MS. **(below)** Extracted ion chromatograms of the reaction products produced by incubation of DacB2 with peptidoglycan substrates.

We next sought to localize a D,D-endopeptidase. As no D,D-endopeptidase has been clearly identified in mycobacteria, we used HHPRED (Zimmermann et al., 2017) to find candidates. By homology to the *E. coli* protein AmpH, an enzyme with both D,D-carboxy- and endopeptidase activity (González-Leiza, de Pedro, & Ayala, 2011), we identified DacB2 (MSMEG_2433), a protein previously shown to have D,D-carboxypeptidase activity in Msm (Bansal et al., 2015), as a candidate to also harbor D,D-endopeptidase capability. We expressed and purified DacB2 and found that it, like AmpH, had both D,D-carboxypeptidase and D,D-endopeptidase activity on peptidoglycan substrates generated *in vitro* (*Figure 4C, Figure 4-figure supplement 1A-C*). Consistent with our model, DacB2-mRFP localized closer to LDT-mRFP, farther from the poles, at sites of older PG (*Figure 4A, B, Figure 4 – video 3*). Thus, bleb formation may result from unchecked D,D-endopeptidase activity in ΔLDT cells.

### Drugs targeting both PBPs and LDTs synergize to kill Mtb

The importance of 3-3 crosslinks in mycobacteria suggests a unique vulnerability. While Mtb can be killed by most non-carbapenem (N-C) b-lactams like penicillins, which largely target the PBPs, carbapenem b-lactams, which target both PBPs and LDTs (Mainardi et al., 2007; Papp-Wallace, Endimiani, Taracila, & Bonomo, 2011) are also effective against Mtb (Diacon et al., 2016; Hugonnet et al., 2009). As has been previously proposed (Gonzalo & Drobniewski, 2013; Gupta et al., 2010; Mainardi et al., 2007), our data suggest that more rapid killing of Mtb could be achieved with drug combinations that target both PBPs and LDTs (*Figure 5A*). In fact, we find that the combination of amoxicillin (a penicillin) and meropenem (a carbapenem), exhibits synergism in minimal inhibitory concentration (S Fractional Inhibitory Concentration < 0.5 (“Synergism Testing: Broth Microdilution Checkerboard and Broth Macrodilution Methods,” 2016), *Figure 5B, Figure 5-figure supplement 1A,B*). We also see that this combination leads to more rapid killing of Mtb *in vitro* (*Figure 5B, Figure 5-figure supplement 2A*) (Andreu, Fletcher, Krishnan, Wiles, & Robertson, 2012).

**Figure 5:**
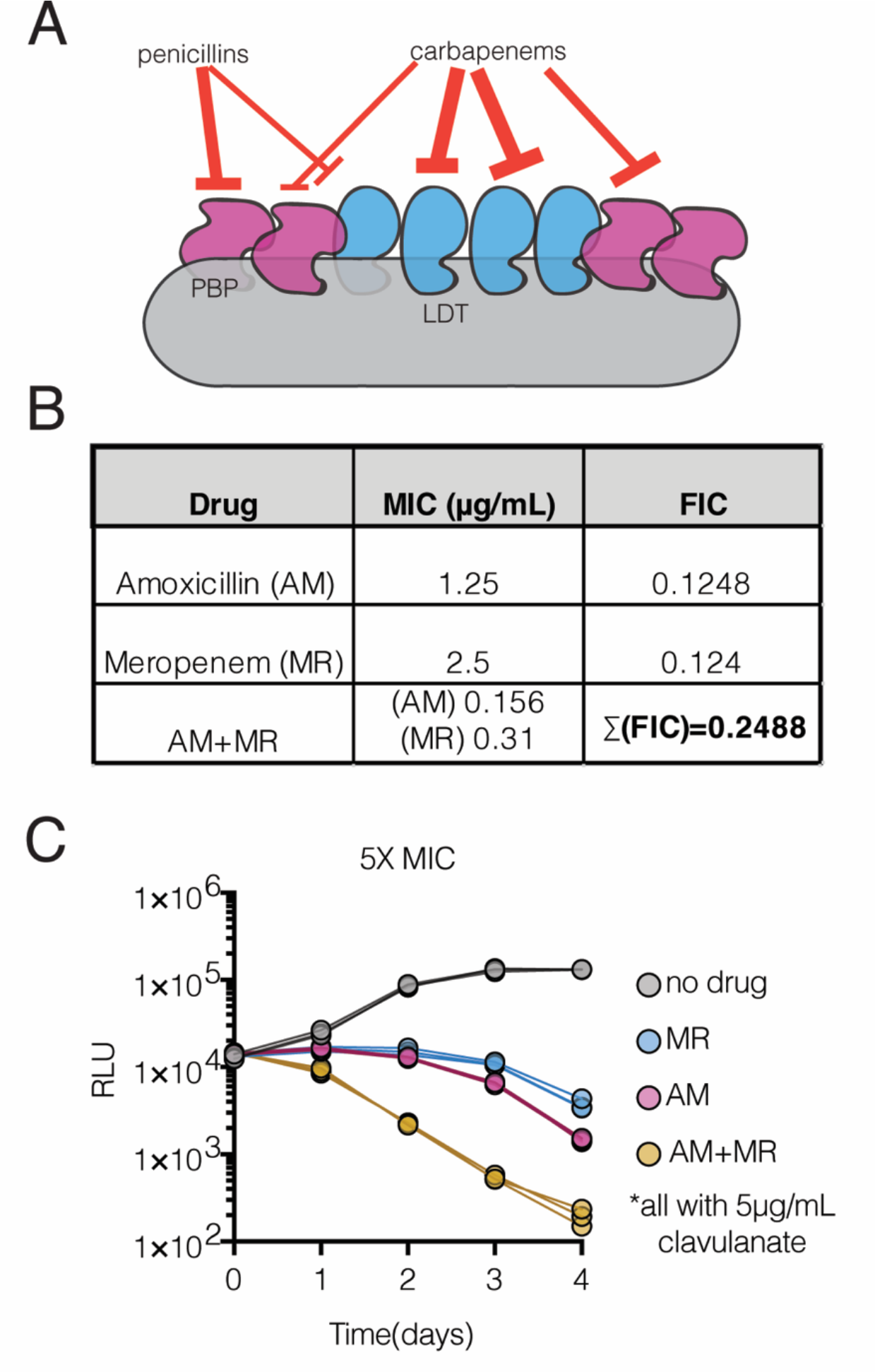
Drugs targeting both PBPs and LDTs synergize to kill Mtb. **(A)** Schematic of synergistic targeting of PBPs and LDTs by pencillin and carbapenem b-lactams. **(B)** Minimum inhibitory concentration (MIC) of amoxicillin, meropenem, or the combination, in *M. tuberculosis*. FIC (fractional inhibitory concentration) = MIC of drug in combination/MIC of drug alone. Synergy is defined as ∑ FIC<=0.5 (“Synergism Testing: Broth Microdilution Checkerboard and Broth Macrodilution Methods,” 2016). **(C)** Killing dynamics of *M. tuberculosis* (expressing the *luxABCDE* operon from *Photorhabdus luminescens* (Andreu et al., 2010)) measured via luciferase production (RLU=relative light units). 5X MIC Amoxicillin (AM) (3.125 µg/mL); 5X MIC Meropenem (MR) (6.25 µg/mL); Amoxicillin+Meropenem 3.125 µg/mL AM; 6.25 µg/mL MR). Biological triplicate are plotted. All drugs were used in combination with 5µg/mL clavulanate.

**Figure 6:**
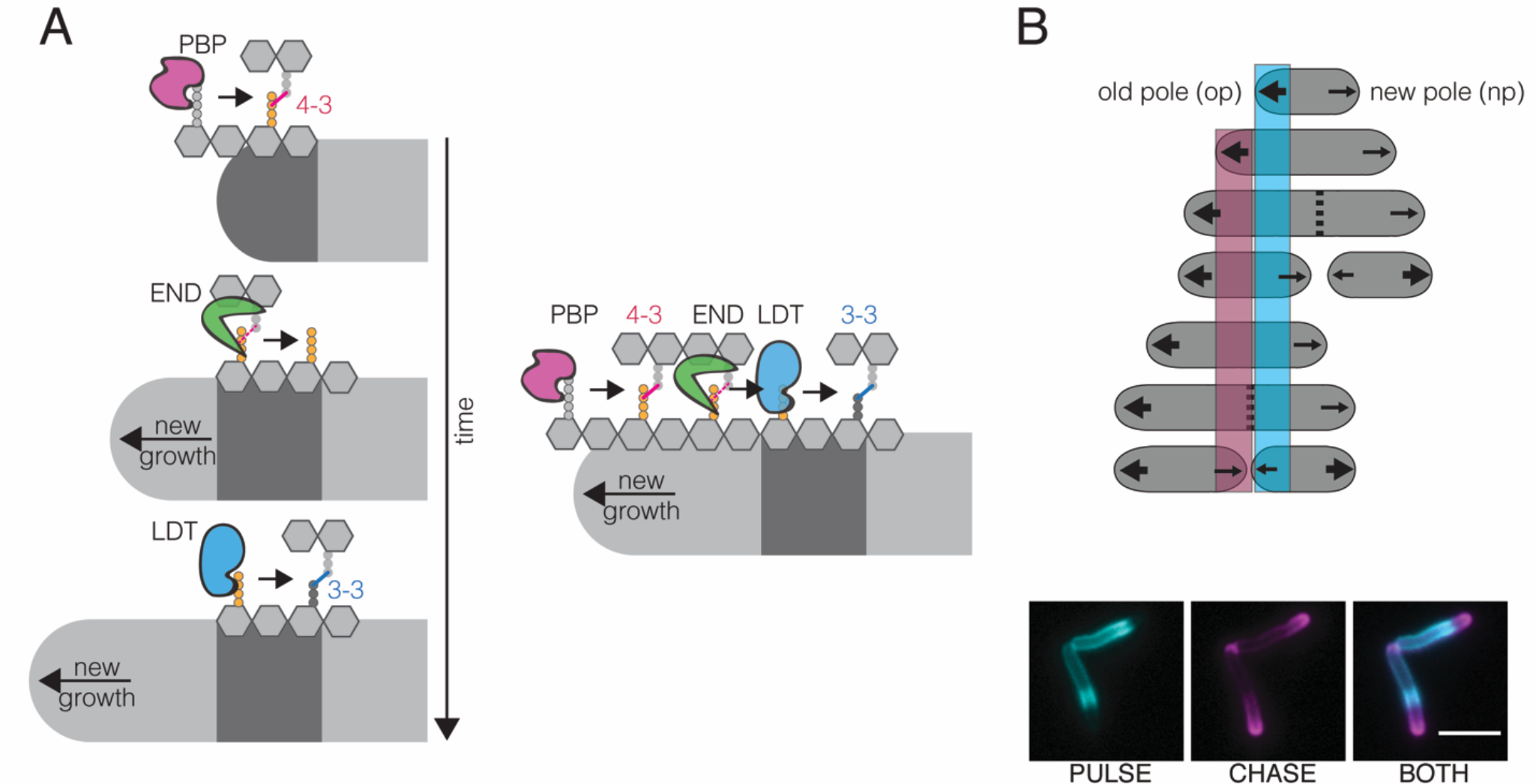
Model for PG enzyme and substrate distribution as governed by polar growth and PG segregation by age. **(A)** A model for PG age, PG enzyme and crosslink segregation via polar growth in mycobacteria. **(B)** Schematic of PG segregation by age (top). 2-minute FDAA pulse (cyan), 45-minute outgrowth, followed by 2-minute FDAA chase (magenta) in WT Msm cells (bottom). Newest cell wall (magenta), older cell wall (cyan). Scale bar=5μm

## Discussion

The success of antibiotics that target PG, like b-lactams, has led to decades of research of this critical bacterial polymer. Recently developed fluorescent probes (FDAAs) of PG synthesis have been used extensively to study PG-synthesis in live cells of numerous bacterial species. Intriguingly, these probes can be incorporated through diverse pathways in different bacteria and thus, their pattern can mark distinct processes (Kuru et al., 2012). Here, we find that, in mycobacteria, FDAA incorporation is primarily LDT-dependent. FDAA incorporation in Msm shows an unusual gradient pattern (Botella et al., 2017), suggesting an asymmetric distribution of tetrapeptide substrate for the LDT-dependent exchange reaction. In addition to their ability to exchange D-amino acids onto tetrapeptides, LDTs also catalyze non-canonical 3-3 crosslinks.

LDT-catalyzed crosslinks are rare in model rod-shaped bacteria like *E.coli* and *B. subtilis* but, are abundant in polar growing bacteria like mycobacteria, *Agrobacterium tumefaciens* and *Sinorhizobium meliloti* (Brown et al., 2012; Cameron, Zupan, & Zambryski, 2015; Kumar et al., 2012; Lavollay et al., 2008; Pisabarro et al., 1985). Here, we find that Msm cells lacking 3-3 crosslinks cannot maintain rod shape at sites of aging cell wall. PBP-catalyzed 4-3 crosslinks appear able to maintain rod shape near the poles, the sites of newer cell wall (*Figure 6A*). Over time, older cell wall moves towards the middle of the cell, loses structural stability, and begins to bleb. The gradual manner in which rod shape is lost suggests that cell wall processing must occur to de-stabilize this portion of the rod. Consistent with this idea, we find that an enzyme that cleaves 4-3 crosslinks, DacB2, also localizes to sites of old cell wall.

Why would Msm cells create 4-3 crosslinks to eventually cleave them? There are many possibilities. For example, perhaps in the absence of lateral cell wall synthesis, the creation of substrate for LDTs through the destruction of 4-3 crosslinks allows the cell to engage the PG along the lateral cell body. This could be important for altering the thickness of the PG layer or anchoring it to the membrane at sites of otherwise “inert” cell wall. Additionally, it may be that as PG ages, it is being manicured for septal synthesis. Supporting this idea, we find that the localization patterns of PonA1, LdtE and DacB2, and FDAA all have local minima at mid-cell closer to the new pole, the asymmetric site of division in mycobacteria (Aldridge et al., 2012; Dhar, Bousbaine, Wakamoto, McKinney, & Santi, 2013; Eskandarian et al., 2017). This lack of localization suggests a lack of penta- and tetra-peptide substrates, implying that that this region of the cell more abundantly crosslinked, as crosslinking utilizes these peptide species. Mycobacteria are missing known molecular septal placement mechanisms like the Noc and the Min system (Hett & Rubin, 2008). Could 3-3 crosslinking be a signal for septal placement? Indeed, the major septal PG hydrolase is RipA, a D,L-endopeptidase which cleaves the bond between the second and third amino acid of PG side chains, a substrate available on 3-3 crosslinked material (Böth, Schneider, & Schnell, 2011; Vollmer, Joris, Charlier, & Foster, 2008b). Thus, it is intriguing to speculate that 3-3 crosslinks could be important for localizing cell division machinery.

In well-studied rod-shaped bacteria like *E.coli* and *B. subtilis*, shape is maintained by MreB-directed PG synthesis along the lateral cell body (Garner et al., 2011; Hussain et al., 2018; Ursell et al., 2014). On the other hand, mycobacteria maintain shape in the absence of an obvious MreB homolog, and in the absence of lateral cell wall elongation. Furthermore, in contrast to lateral-elongating bacteria, in which new and old cell wall are constantly intermingled during growth, polar growth segregates new and old cell wall (*Figure 6B*). We find that mycobacteria appear to utilize 3-3 crosslinks at asymmetrically distributed aging cell wall to provide stability along the lateral body, something that may not be required in the presence of MreB-directed PG synthesis.

New drug combinations for TB are desperately needed. Indeed, there has been a renewed interest in repurposing FDA-approved drugs for TB treatment (Diacon et al., 2016). Our results suggest that a better understanding of the localization and distribution of drug targets in single cells may lead to rational predictions of drug combinations. Indeed, we find that enzymes which do very similar chemistry – PG crosslinking – are distributed differentially in a single cell and across the population. In the absence of 3-3 crosslinks, 4-3 crosslinks become more important for cell viability. These data predict that a drug combination which inhibits both PBPs and LDTs will work synergistically to more quickly kill Mtb, a prediction we verified *in vitro*. Interestingly, meropenem combined with amoxicillin/clavulanate resulted in early clearance of Mtb from patient sputum (Diacon et al., 2016). In fact, the combination might be key to accelerated killing of Mtb (Gonzalo & Drobniewski, 2013).

## Acknowledgments

We thank Cara Boutte and Erkin Kuru for discussion and thorough manuscript review. We are grateful to the Rubin and Fortune laboratory for discussion and input. We also thank the Microbiology and Immunobiology department at Harvard Medical School for sharing equipment and reagents, as well as the BSL3 staff at the Harvard School of Public health for tremendous support. This work was supported by the National Institutes of Health U19 AI107774 to E.J.R and T.R.I., F32AI104287 to E.H.R (who is also supported by a Career Award at the Scientific Interface from BWF), R01 GM76710 to S.W., R01AI083365 and U19AI109764 to T.G.B.. M.A.W. is supported by an F32 GM123579. LT.S. was supported by an American Heart Association Postdoctoral fellowship (14POST18480014). H.C.L is funded by a Simons Foundation Fellow of the Life Sciences Research Foundation award. This work was also supported in part by the Swiss National Science Foundation (310030_156945) and the Innovative Medicines Initiative (115337) to J.D.M., and from the Swiss National Science Foundation (205321_134786 and 205320_152675), the European Union FP7/2007-2013/ERC Grant Agreement No. 307338 (NaMic), and EU-FP7/Eurostars E!8213 to G.E.F. Support for H.A.E. comes from a European Molecular Biology Organization Long Term Fellowships (191-2014 and 750-2016). K.J.K. was supported by the National Science Foundation Graduate Research Fellowship (DGE1144152, DGE0946799).

## Materials and Methods

### Bacterial strains and culture conditions

Unless otherwise stated, *Mycobacterium smegmatis* (mc^2^155) was grown shaking at 37°C in liquid 7H9 media consisting of Middlebrook 7H9 salts with 0.2% glycerol, 0.85g/L NaCl, ADC (5g/L albumin, 2g/L dextrose, 0.003g/L catalase), and 0.05% Tween 80 and plated on LB agar. *Mycobacterium tuberculosis* (H37Rv) was grown in liquid 7H9 with OADC (oleic acid, albumin, dextrose, catalase) with 0.2% glycerol and 0.05% Tween 80. Antibiotic selection for *M. smegmatis* and *M. tuberculosis* were done at the following concentrations in broth and on agar: 25µg/mL kanamycin, 50µg/mL hygromycin, 20µg/mL zeocin and 20µg/mL nourseothricin and, twice those concentrations for cloning in *Escherichia coli* (TOP10, XL1-Blue and DH5α).

### Strain construction

*M. smegmatis* mc^2^155 mutants lacking *ldtABECFG* (DLDT) was constructed using recombineering to replace endogenous copies with zeocin or hygromycin resistance cassettes flanked by lox sites as previously described (Boutte et al., 2016). Briefly, 500 base pairs of upstream and downstream sequence surrounding the gene of interest were amplified via PCR (KOD XtremeTM Hot Start DNA polymerase (EMD Millipore, Billerica, MA)). These flanking regions were amplified with overlaps to either a zeocin or hygromycin resistance cassette flanked by loxP sites and these pieces were assembled into a deletion construct via isothermal assembly (Gibson et al., 2009). Each deletion cassette was transformed into Msm expressing inducible copies of RecET for recombination (Murphy, Papavinasasundaram, & Sassetti, 2015). Once deletions were verified by PCR using and sequencing, the antibiotic resistance cassettes were removed by the expression of Cre recombinase. The order of deletion construction in the ΔLDT strain was as follows (where arrows represent transformation of a Crerecombinase plasmid, followed by curing of the Cre-recombinase plasmid as it contains the *sacB* gene for sucrose counter selection on LB supplemented with 10% sucrose, and strain names are listed in parenthesis). This resulted in the removal of antibiotic cassettes flanked by loxP sites:

1. mc^2^155*ΔldtA*:: zeo^R^ (KB103)à mc^2^155*ΔldtA*::loxP (KB134)
2. mc^2^155*ΔldtA::loxP +* Δ*ldtE*:: zeo^R^ (KB156)
3. mc^2^155*ΔldtA::loxP* Δ*ldtE*:: zeo^R^ + Δ*ldtB*:: hyg^R^ (KB200) à mc^2^155*ΔldtA::loxP* Δ*ldtE*::loxP Δ*ldtB*::loxP (KB207)
4. mc^2^155*ΔldtA::loxP* Δ*ldtE*::loxP Δ*ldtB*::loxP + Δ*ldtC*:: hyg^R^ (KB209)
5. mc^2^155*ΔldtA::loxP* Δ*ldtE*::loxP Δ*ldtB*::loxP Δ*ldtC*:: hyg^R^ Δ*ldtG*:: zeo^R^ (KB222)à mc^2^155*ΔldtA::*loxP Δ*ldtE*::loxP Δ*ldtB*::loxP Δ*ldtC*:: loxP Δ*ldtG*:: loxP (KB241)
6. mc^2^155*ΔldtA::*loxP Δ*ldtE*::loxP Δ*ldtB*::loxP Δ*ldtC*:: loxP Δ*ldtG*:: loxP Δ*ldtF*:: hyg^R^ (KB303 referred to as DLDT).

*M. tuberculosis* H37Rv was transformed with a vector expressing the codon optimized *Photorhabdus luminescens luxABCDE* operon (pMV306hsp+LuxG13 –Addgene #26161)(Andreu et al., 2010). This strain is referred to as Mtb Lux.

Refer to Supplemental Table 1 for oligonucleotides, and Supplemental Table 2 for a complete strain list.

### ΔLDT complementation plasmid construction

To complement ΔLDT we placed a copy of *ldtE* (*MSMEG_0233*) under the constitutive TetO promoter (a UV15 derivative within a pMC1s plasmid that is inducible with anhydrous tetracycline in the presence of a tet-repressor TetR, which the ΔLDT_comp_ strain lacks (Kieser, Boutte, et al., 2015b)) on vector that integrates at the L5 phage integration site of the chromosome of the ΔLDT strain (the vector is marked with kanamycin resistance). A glycine, glycine, serine linker was cloned between *ldtE* and mRFP in this complementation construct.

### PonA1 transpeptidase essentiality L5 allele swapping

To test essentiality of transpeptidation by PonA1 in the ΔLDT cells, L5 allele swapping as described in (Kieser, Boutte, et al., 2015b) was performed. The plasmids used in this experiment were previously published in (Kieser, Boutte, et al., 2015b). Briefly, a wild-type copy of PonA1 (TetO driven expression, L5 integrating and kanamycin marked) was transformed into ΔLDT. Then, the endogenous copy of *ponA1* was replaced with zeocin using the above mentioned recombineering technique (amplifying the construct from a previously published deletion mutation of *ponA1*(Kieser, Boutte, et al., 2015b)). Swapping efficiency of either wildtype or transpeptidase mutant PonA1 marked with nourseothricin was tested with a transformation into ΔLDT//L5-TetO-ponA1 (WT)-kanamycin.

### *M. tuberculosis* minimum inhibitory concentration (MIC) determination

Mtb Lux was grown to log phase and diluted to an OD_600_=0.006 in each well of non-treated 96 well plates (Genessee Scientific) containing 100µL of meropenem (Sigma Aldrich) and/or amoxicillin (Sigma Aldrich) diluted in 7H9+OADC+5µg/mL clavulanate (Sigma Aldrich). Cells were incubated in drug at 37°C shaking for 7 days, 0.002% resazurin (Sigma Aldrich) was added to each well, and the plates were incubated for 24 hours before MICs were determined. Pink wells signify metabolic activity and blue signify no metabolic activity. (Kieser, Baranowski, et al., 2015a) Checkerboard MIC plates and fractional inhibitory concentrations were calculated as described in (“Synergism Testing: Broth Microdilution Checkerboard and Broth Macrodilution Methods,” 2016).

### *M. tuberculosis* drug killing assays

Mtb Lux was grown to log phase (kanamycin 25µg/mL) and diluted in 30mL inkwells (Corning Lifesciences) to an OD_600_=0.05 in 7H9+OADC+5µg/mL clavulanate with varying concentrations of amoxicillin, meropenem, or both. 100µL of these cultures were pipetted in duplicate into a white 96-well polystyrene plate (Greiner Bio-One) and luminescence was read in a Synergy H1 microplate reader from BioTek Instrumenmts, Inc. using the Gen5 Software (2.02.11 Installation version). The correlation between relative light units (RLU) and colony forming units (CFU) is shown in Msm in Figure 5 – figure supplement 2.

### Fluorescent D-amino acid labeling

NADA (3-[7-nitrobenzofurazan]-carboxamide-D-alanine), HADA (3-[7-hydroxycoumarin]-carboxamide-D-alanine) and TADA (3-[5-carboxytetramethylrhodamine]-carboxamide-D-alanine) were synthesized by Tocris following the published procotol (Kuru, Tekkam, Hall, Brun, & Van Nieuwenhze, 2014). To 1mL of exponentially growing cells 0.1mM of FDAA final was added and incubated for 2 minutes before washing in 7H9 twice. For still imaging, after the second wash, cells were fixed in 7H9 + 1% paraformaldehyde before imaging. For pulse chase experiments, cells were stained, washed with 7H9 and allowed to grow out for 40 minutes before being stained with a second dye and imaged.

### Flow cytometry

An *M. smegmatis* transposon library was grown to mid-log phase, and stained with 2 µg/mL NADA for 2 minutes. Cells were centrifuged and half of the supernatant was discarded. The pellet was resuspended in the remaining supernatant, passed through a 10µm filter and taken to be sorted (FACSAria; Excitation: 488nm; Emission filter: 530/30). Two bins were drawn at the lowest and highest staining end of the population, representing 12.5% of the population. 600,000 cells from these bins were sorted into 7H9 medium. Half of this was directly plated onto LB agar supplemented with kanamycin to select for cells harboring the transposon. The remaining 300,000 cells were grown out in 7H9 to log phase, stained with FDAA and sorted again to enrich the populations.

### Transposon sequencing, mapping and FDAA flow cytometry enrichment analysis

Genomic DNA (gDNA) was harvested from the sorted transposon library colonies and transposon-gDNA junction libraries were constructed and sequenced using the Illumina Hi-Seq platform (Long et al., 2015). Reads were mapped on the *M. smegmatis* genome, tallied and reads at each TA site for the bins (low/high incorporating sort 1 and 2) were imported into MATLAB and processed by a custom scripts as described in (Rego, Audette, & Rubin, 2017).

### Microscopy

Both still imaging and time lapse microscopy were performed on an inverted Nikon TI-E microscope at 60x magnification. Time lapse was done using a CellASIC (B04A plate) with constant liquid 7H9 flow in a 37°C chamber. For turgor experiment (*Figure 2A*), cells were grown in either 7H9 or 7H9 500mM sorbitol overnight, and then switched to either 7H9 with 150mM sorbitol (high osmolar) or to 7H9 alone (iso-osmolar).

### Atomic force microscopy

AFM experimentation was conducted as previously (Eskandarian et al., 2017). In short, polydimethylsiloxane (PDMS) – coated coverslips were prepared by spin-coating a mixture of PDMS at a ratio of 15:1 (elastomer:curing agent) with hexane (Sigma 296090) at a ratio of 1:10 (PDMS:hexane) (Koschwanez, Carlson, & Meldrum, 2009; Thangawng, Ruoff, Swartz, & Glucksberg, 2007). A 50 µl filtered (0.5 µm pore size PVDF filter – Millipore) aliquot of bacteria grown to mid-exponential phase and concentrated from 2-5 ml of culture was deposited onto the hydrophobic surface of a PDMS-coated coverslip and incubated for ~20 min to increase surface interactions between bacteria and the coverslip. 7H9 medium (~3 ml) was supplied to the sample so as to immerse the bacterial sample and the AFM cantilever in fluid. The AFM imaging mode, Peak Force QNM, was used to image bacteria with a Nanoscope 5 controller (Veeco Metrology) at a scan rate of 0.5 Hz and a maximum Z-range of 12 µm. A ScanAsyst fluid cantilever (Bruker) was used. Height, peak force error, DMT modulus, and log DMT modulus were recorded for all scanned images in the trace and retrace directions. Images were processed using Gwyddion (Department of Nanometrology, Czech Metrology Institute). ImageJ was used for extracting bacterial cell profiles in a tabular form.

### Correlated optical fluorescence and AFM

Correlated optical fluorescence and AFM images were acquired as described (Eskandarian et al., 2017). Briefly, optical fluorescence images were acquired with an electron-multiplying charge-coupled device (EMCCD) iXon Ultra 897 camera (Andor) mounted on an IX81 inverted optical microscope (Olympus) equipped with an UPLFLN100XO2PH x100 oil immersion objective (Olympus). Transmitted light illumination was provided by a 12V/100W AHS-LAMP halogen lamp. An U-MGFPHQ fluorescence filter cube for GFP with HQ-Ion-coated filters was used to detect GFP fluorescence. The AFM was mounted on top of the inverted microscope, and images were acquired with a Dimension Icon scan head (Bruker) using ScanAsyst fluid cantilevers (Bruker) with a nominal spring constant of 0.7 N m^-^1 in Peak Force QNM mode at a force setpoint ~1nN and typical scan rates of 0.5 Hz. Indentation on the cell surface was estimated to be ~10 nm in the Z-axis. Optical fluorescence microscopy was used to identify Wag31-GFP puncta expressed in a wild-type background(Dhar et al., 2013) in order to distinguish them from cells of the ∆LDT mutant strains.

### Calculating cell surface rigidity

A cell profile was extracted from AFM Height and DMT Modulus image channels as sequentially connected linear segments following the midline of an individual cell. A background correction was conducted to by dividing the DMT modulus values of the cell surface by the mean value of the PDMS surface and rescaled to compare the cell surface rigidity between individual cells from different experiments. The DMT modulus reflects the elastic modulus (stress-strain relationship) for each cross-sectional increment along the cell length.

### Analysis of fluorescent protein distribution

Using a segmented line, profiles of cells from new to old pole were created at the frame “pre-division” based on physical cell separation of the phase image. A custom FIJI (Schindelin et al., 2012) script was run to extract fluorescence line profiles of each cell and save them as .csv files. These .csv files were imported to Matlab where a custom script was applied to normalize the fluorescence line profile to fractional cell length and to interpolate the fluorescence values to allow for averaging.

### Analysis of cell wall distribution

Cells were stained with Alexa488 NHS ester as described previously (Aldridge et al., 2012) and followed via time-lapse microscopy in the CellASIC device. Briefly, 1mL of log phase cells was pelleted at 8,000 rpm for 1 minute and washed with 1mL PBST. The pellet was resuspended in 100uL of PBST and 10uL Alexa Fluor 488 carboxylic acid succinimidyl ester was added for a final concentration of 0.05mg/mL. This was incubated for 3 minutes at room temperature. Stained cells were pelleted for 1 minute at 13,000 rpm and washed with 500µL PBST. They were spun again and resuspended in 7H9 for outgrowth observation over time in the CellASIC device.

### Analysis of FDAAs

Images were analyzed using a combination of Oufti (Paintdakhi et al., 2016) for cell selection followed by custom coded Matlab scripts to plot FDAA fluorescence over normalized cell length, calculate cell length and bin cells by existence of an FDAA labeled septum.

### Generation of transposon libraries

*M. smegmatis* cells were transduced at (OD_600_ 1.1-1.7) with *φ*MycoMarT7 phage (temperature sensitive) that has a Kanamycin marked Mariner transposon as previously described (Long et al., 2015). Briefly, mutagenized cells were plated at 37°C on LB plates supplemented with Kanamycin to select for phage transduced cells. Roughly 100,000 colonies per library were scraped, and genomic DNA was extracted. Sequencing libraries were generated specifically containing transposon disrupted DNA. Libraries were sequenced on the Illumina platform. Data were analyzed using the TRANSIT pipeline (DeJesus et al., 2015).

### Peptidoglycan isolation and analysis

600mLs of wildtype and ΔLDT cells were grown to log phase and collected via centrifugation at 5000 x g for 10 minutes at 4°C. The resulting pellet was resuspsended in PBS and cells were lysed using a cell disruptor at 35,000psi twice. Lysed cells were boiled in 10% SDS (sodium dodecyl sulfate) for 30 minutes and peptidoglycan was collected via centrifugation at 17,000 x g. Pellets were washed with 0.01% DDM(*n-*Dodecyl β-D-maltoside) to remove SDS and resuspended in 1XPBS + 0.01% DDM. PG was digested with alpha amylase (Sigma A-6380) and alpha chymotrypsin (Amersco 0164) overnight. The samples were again boiled in 10% SDS and washed in 0.01% DDM. The resulting pellet was resuspended in 400µL 25mM sodium phosphate pH6, 0.5mM MgCl2, 0.01% DDM. 20µL of lysozyme (10mg/mL) and 20µL 5U/µL mutanolysoin (Sigma M9901) were added and incubated overnight at 37°C. Samples were heated at 100°C and centrifuged at 100,000 x g. 128µL of ammonium hydroxide was added and incubated for 5 hours at 37°C. This reaction was neutralized with 122µL of glacial acetic acid. Samples were lyophilized, resuspended in 300µL 0.1% formic acid and subjected to analysis by LC-MS/MS. Peptide fragments were separated with an Agilent Technologies 1200 series HPLC on a Nucleosil C18 column (5µm 100A 4.6 x 250mm) at 0.5mL/min flow rate with the following method: Buffer A= 0.1% Formic Acid; Buffer B=0.1% Formic Acid in acetonitrile; 0% B from 0-10 minutes, 0-20% B from 10-100 minutes, 20% B from 100-120 minutes, 20-80% B from 120-130 minutes, 80% B from 130-140 minutes, 80%-0% B from 140-150minutes, 0% B from 150-170 minutes. MS/MS was conducted in positive ion mode using electrospray ionization on an Agilent Q-TOF (6520).

### Expression and purification of MSMEG_2433 (DacB2)

MSMEG_2433 was expressed and purified using a modified method for purification of low molecular weight PBPs that was previously published (Qiao et al., 2014). An N-terminally truncated MSMEG_2433_(29-296)_ was cloned into the pET28b vector for isopropyl β-D-1-thiogalactopyranoside (IPTG) inducible expression in *E. coli* BL21 (DE3). 10mLs of overnight culture grown in LB with Kanamycin (50µg/mL) were diluted 1:100 into 1 L of LB with Kanamycin (50µg/mL) and grown at 37°C until an OD_600_ of 0.5. The culture was cooled to room temperature, induced with 0.5mM IPTG, and shaken at 16°C overnight. Cells were pelleted via centrifugation at 4,000rpm for 20 min at 4°C. The pellet was suspended in 20mL binding buffer (20mM Tris pH 8, 10mM MgCl_2_, 160mM NaCl, 20mM imidazole) with 1mM phenylmethylsulfonylfluoride (PMSF) and 500μg/mL DNase. Cells were lysed via three passage through a cell disrupter at ≥ 10,000 psi. Lysate was pelleted by ultracentrifugation (90,000 × g, 30 min, 4°C). To the supernatant, 1.0mL washed Ni-NTA resin (Qiagen) was added and the mixture rocked at 4°C for 40 min. After loading onto a gravity column, the resin was washed twice with 10mL wash buffer (20mM Tris pH 8, 500mM NaCl, 20mM imidazole, 0.1% Triton X-100). The protein was eluted in 10mL of elution buffer (20 mM Tris pH8, 150mM NaCl, 300mM imidazole, 0.1% reduced Triton X-100) and was concentrated to 1 mL with a 10kD MWCO Amicon Ultra Centrifuge Filter. The final protein concentration was measured by reading absorbance at 280 nm and using the estimated extinction coefficient (29459 M^−^1cm^−^1) calculate concentration. The protein was diluted to 200μM in elution buffer with 10% glycerol, aliquoted, and stored at −80°C.

Proper folding of purified MSMEG_2433_(29-296)_ was tested via Bocillin-FL binding. Briefly, 20µM of purified protein was added to penicillin G (100, 1000 U/mL in 20mM K_2_HPO_4_, 140mM NaCl, pH7.5) in a 9µL reaction. The reaction was incubated at 37°C for 1hour. 10µM Bocillin-FL was added and incubated at 37°C for 30 minutes. SDS loading dye was added the quench the reaction and samples were loaded onto a 4-20% gel. MSMEG_2433_(29-_296) bound by Bocillin-FL was imaged using a Typhoon 9400 Variable Mode Imager (GE Healthcare) (Alexa Excitation-488nm Emission-526nm).

### Lipid II extraction

*B. subtilis* Lipid II was extracted as previously published (Qiao et al., 2017).

### SgtB purification

*S. aureus* SgtB was purified as previously published (Rebets et al., 2014).

### Purification of B. subtilis PBP1

Purification of *B. subtilis* PBP1 was carried out as previously described (Lebar et al., 2014).

### *In vitro* Lipid II polymerization and crosslinking

20 µM purified BS Lipid II was incubated in reaction buffer (50 mM HEPES pH 7.5, 10 mM CaCl_2_) with either 5 µM PBP1 or 0.33 µM SgtB for 1 hour at room temperature. The enzymes were heat denatured at 95°C for 5 minutes. Purified MSMEG_2433_(29-296)_ was added (20 uM, final) and the reaction was incubated at room temperature for 1 hour. Mutanolysin (1 µL of a 4000 U/mL stock) was added and incubated for 1.5 hours at 37 °C (twice). The resulting muropeptides were reduced with 30 µL of NaBH_4_ (10 mg/mL) for 20 minutes at room temperature with tube flicking every 5 minutes to mix. The pH was adjusted to ~4 using with 20% H_3_PO_4_ and the resulting product was lyophilized to dryness. The residue was resuspended in 18µL of water and analyzed via LC-MS as previously reported (Welsh et al., 2017).

---

**Figure 1- figure supplement 1:**
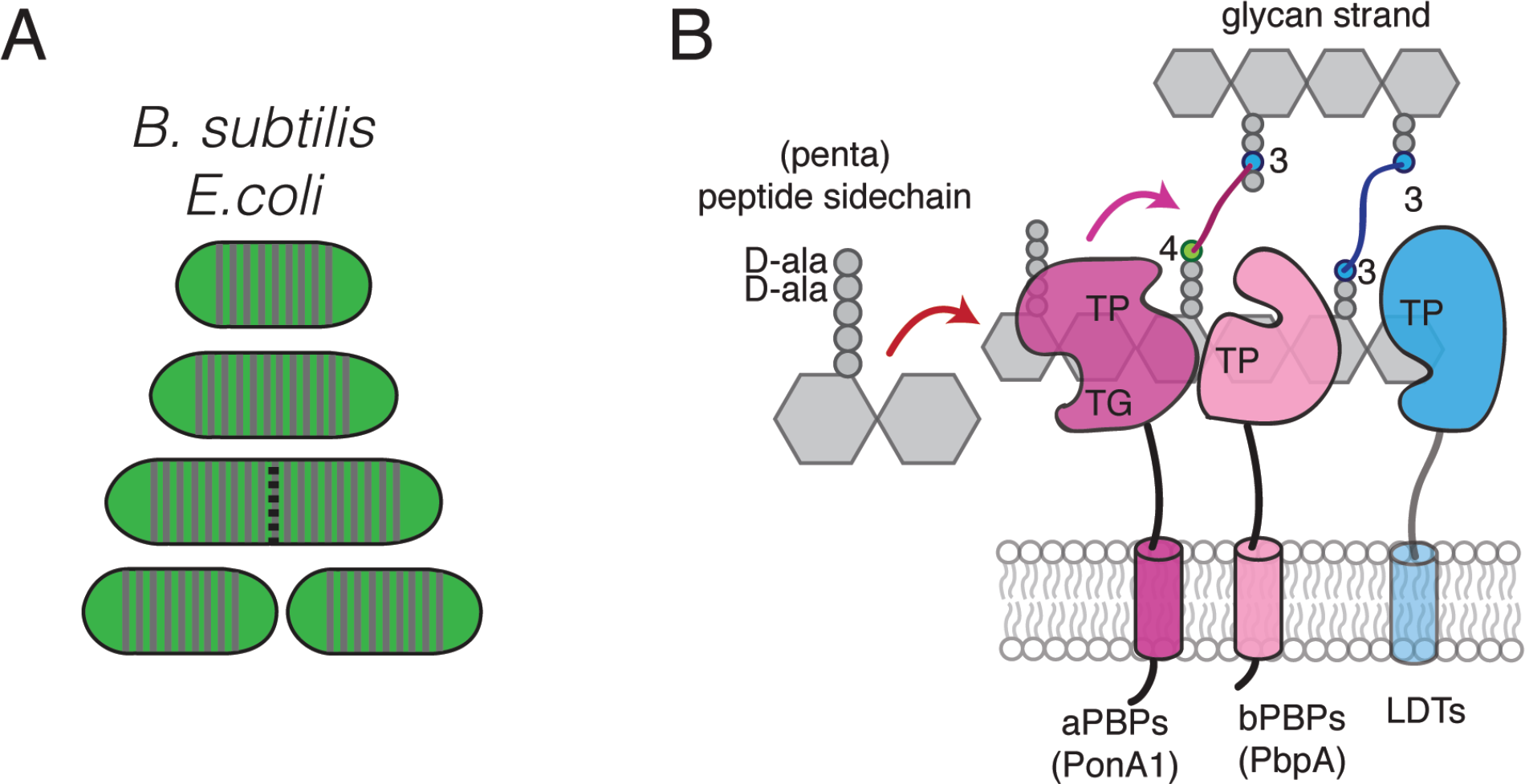
Peptidoglycan synthesis and FDAA screen overview. **(A)** *Escherichia coli* and *Bacillus subtilis* lateral cell wall growth. Unlike mycobacteria, *E. coli* and *B. subtilis* insert new cell wall along the lateral cell body, mixing old and new peptidoglycan. Green portion represents old cell wall; grey portion represents new material. **(B)** Cartoon of penicillin binding proteins (PBPs), L,D-transpeptidases (LDTs), and both 4-3 and 3-3 crosslinking. PBPs utilize a pentapeptide substrate found on new peptidoglycan, ending in D-alanine-D-alanine. LDTs utilize a tetrapeptide substrate found on processed peptidoglycan. TP, transpeptidase; TG, transglycosylase

**Figure 1- figure supplement 2:**
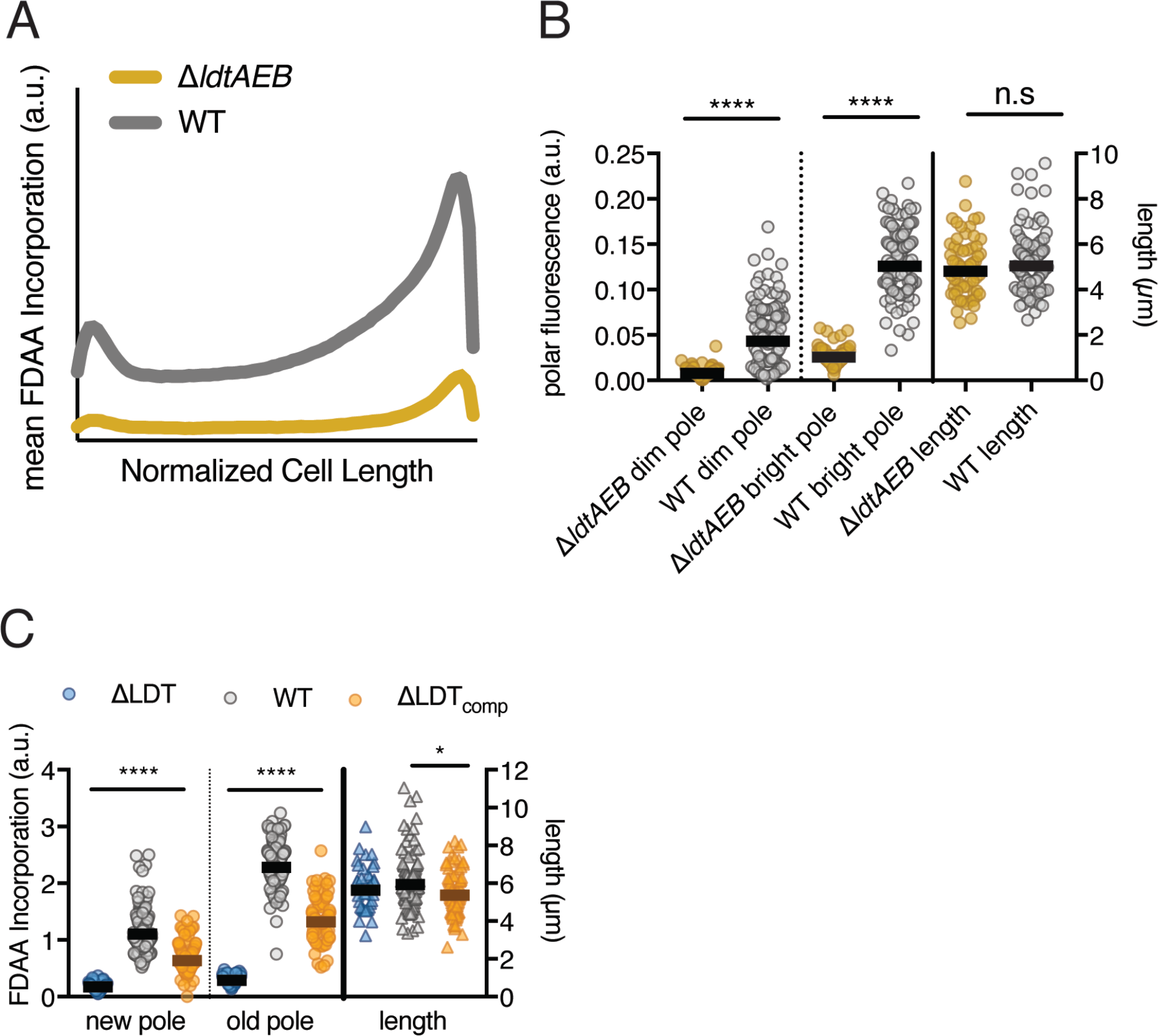
Fluorescent D-amino acid screen validation. **(A)**Mean line profiles (from new to old pole) of FDAA incorporation in log-phase WT (N=97), ∆*ΔldtABE* (N=64). **(B)** Quantification of FDAA incorporation at cell poles and quantification of cell length. Mann-Whitney U P-Value shown (**** P-Value < 0.0001). **(C)** Quantification of FDAA incorporation at poles and cell lengths of WT, ∆LDT and ∆LDT_comp_ cells shown in Fig. 1E and whose mean incorporation is shown in Fig. 1F.

**Figure 1- figure supplement 3:**
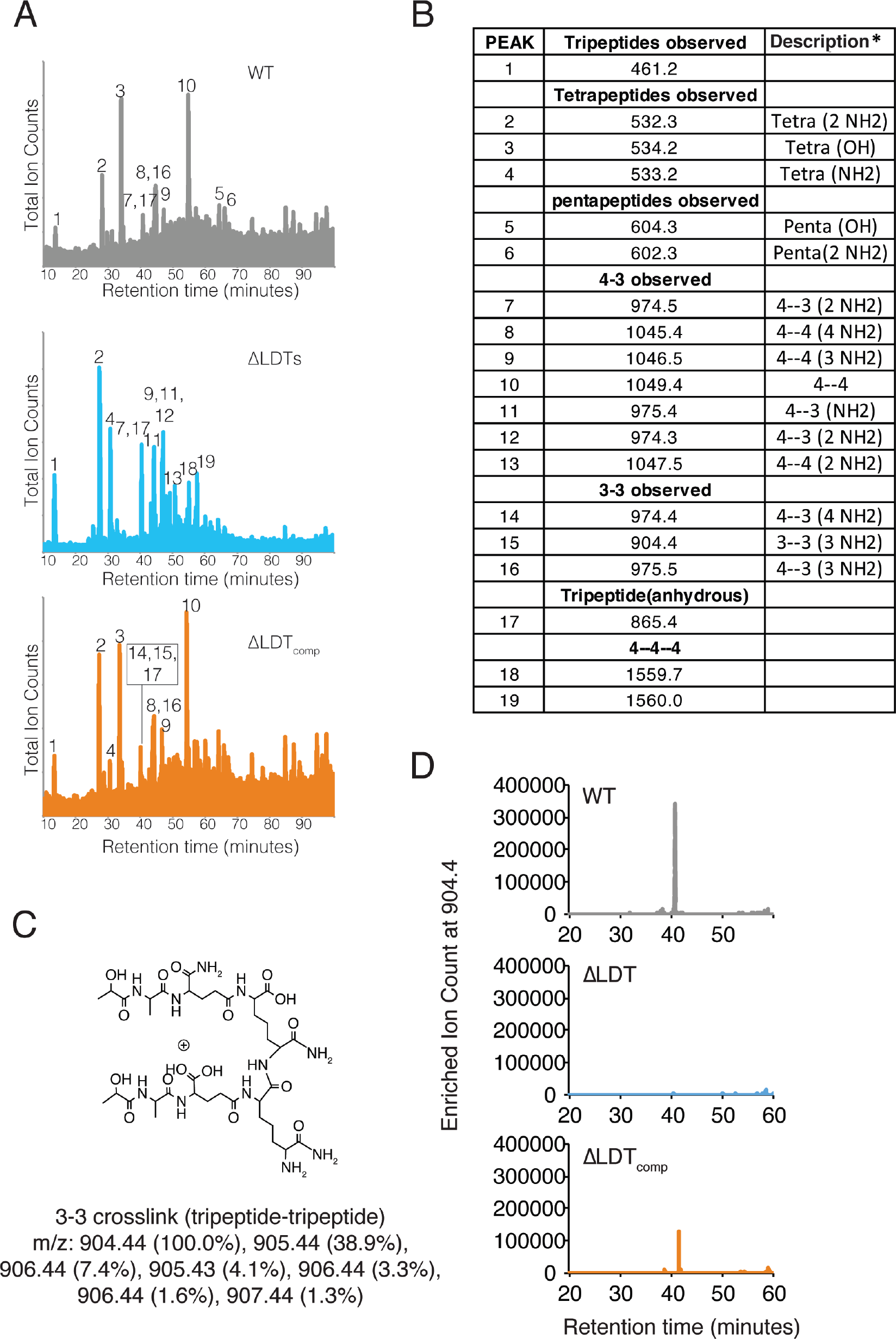
3-3 crosslinks are not detectable in ∆LDT cells. **(A)** Total ion chromatograms of WT, ∆LDT and ∆LDT_comp_ peptidoglycan. **(B)** Table of muropeptide masses (Da) observed in (S3A). The molecular weight difference by one of the identified peptides is due to differential amidation. The descriptions include the peptide lengths in the crosslink (4= tetra-, 3= tri- peptide) and the following parenthesis specifies the number of amidation in the species according to mass. **(C)** Structure of a representative 3-3 crosslink with a m/z=904.4. **(D)** Extracted ion chromatograms from WT, ∆LDT and ∆LDT_comp_ for a representative 3-3 crosslink with a m/z=904.4.

**Figure 2- figure supplement 1:**
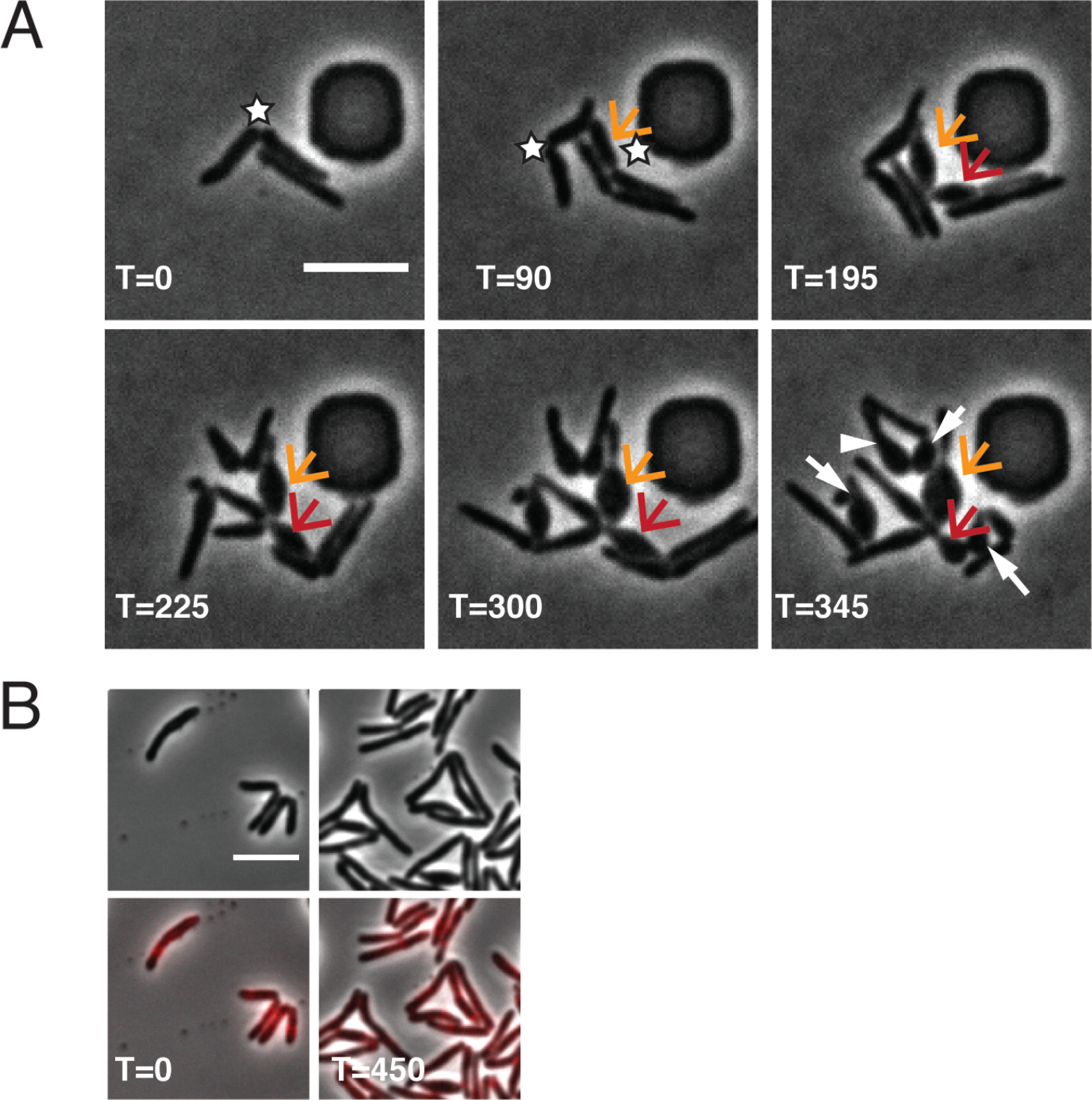
∆LDT cell morphological characteristics. **(A)** Time-lapse microscopy montage of ∆LDT cells. The white stars mark new poles. The orange arrow points to the first new pole daughter cell of this series. The red arrow indicates the second resulting new pole daughter cell. In the last frame, white arrows point to all new pole daughter cells (besides the orange arrow and red arrow). **(B)** Time-lapse microscopy montage of ∆LDT_comp_ cells expressing LdtE-mRFP.

All scale bars=5μm

**Figure 2- figure supplement 2:**
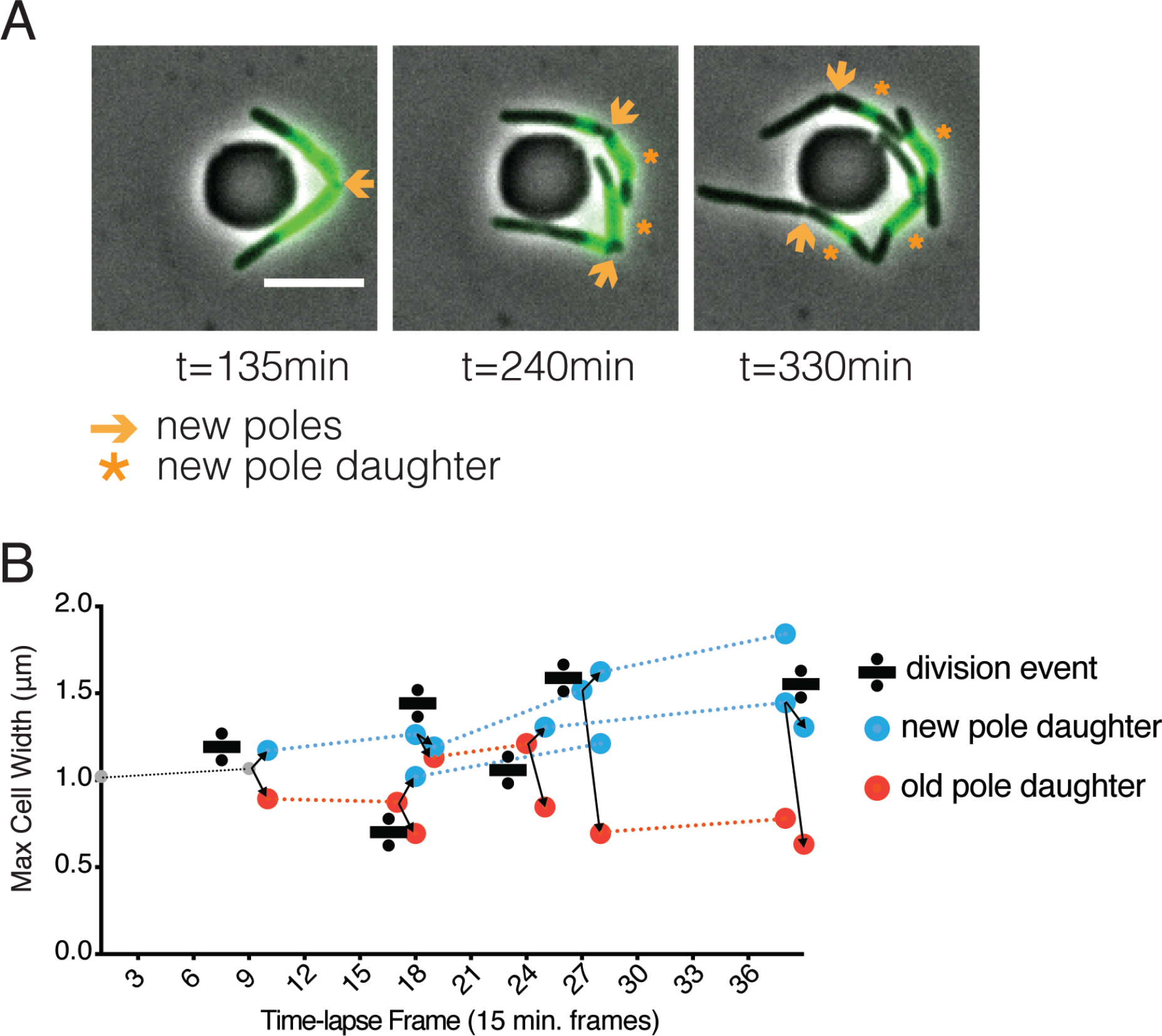
Inheritance of old cell wall and occurrence of blebs in new pole daughter cells. **(A)** WT Msm stained with Alexa Fluor^TM^ 488 NHS ester, washed and visualized over time. New material is unstained, old material is stained green. Orange arrows indicate a new pole. Orange stars mark new pole daughter cells. All scale bars=5μm **(B)** Maximum cell width of ∆LDT cell lineages over time. Width of new pole daughters = blue circle; width of old pole daughters = orange circle. Division signs denote a division event. At each division, there are two arrows from the dividing cell leading to the resulting new and old pole daughter cell widths (blue and orange respectively).

**Figure 3- figure supplement 1:**
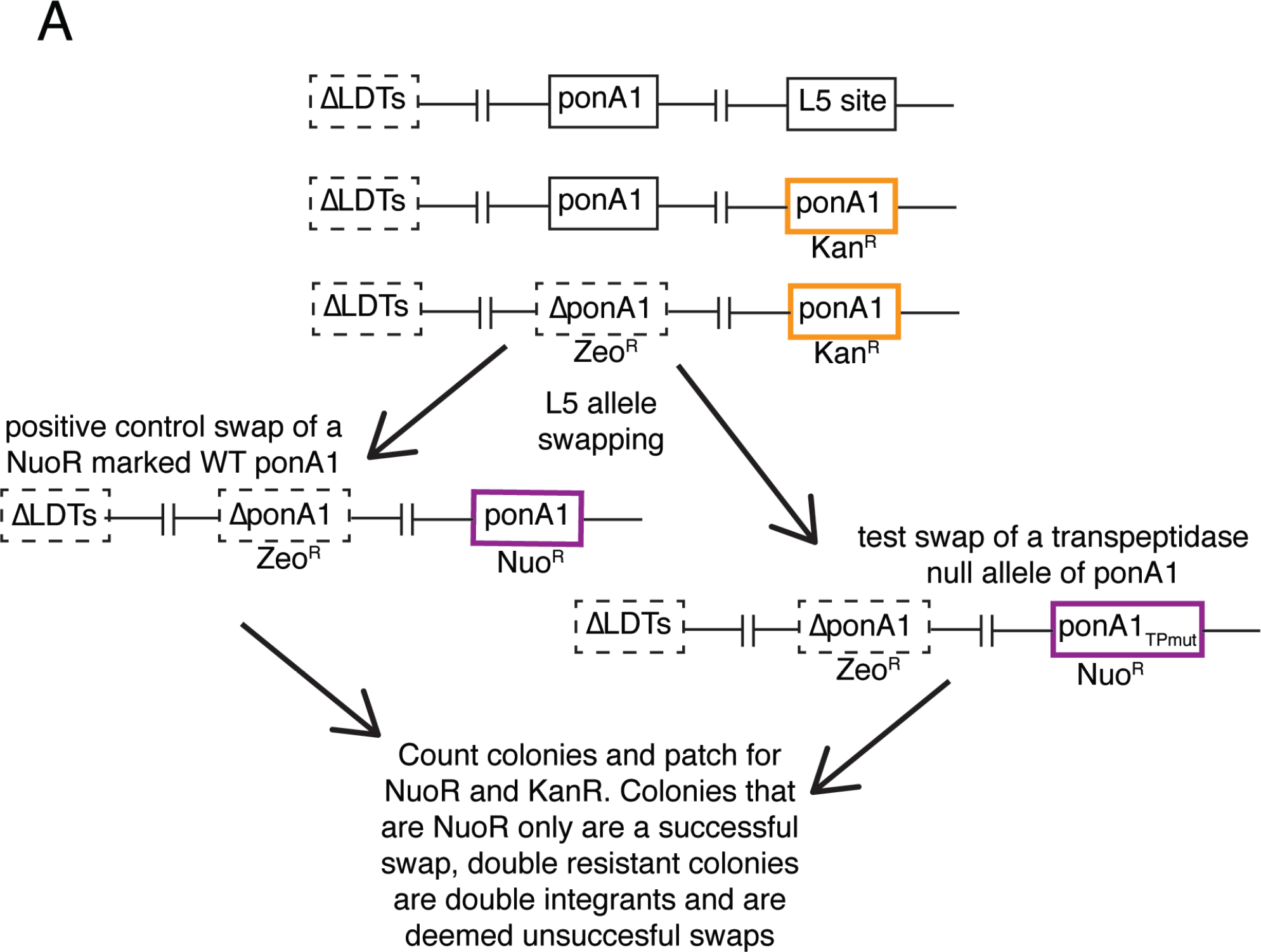
L5 allele swapping to test essentiality of PonA1’s ability to form 4-3 crosslinks (transpeptidation). **(A)** Schematic of L5 allele swapping experiment. Adapted from (Kieser, Boutte, et al., 2015b).

**Figure 4- figure supplement 1:**
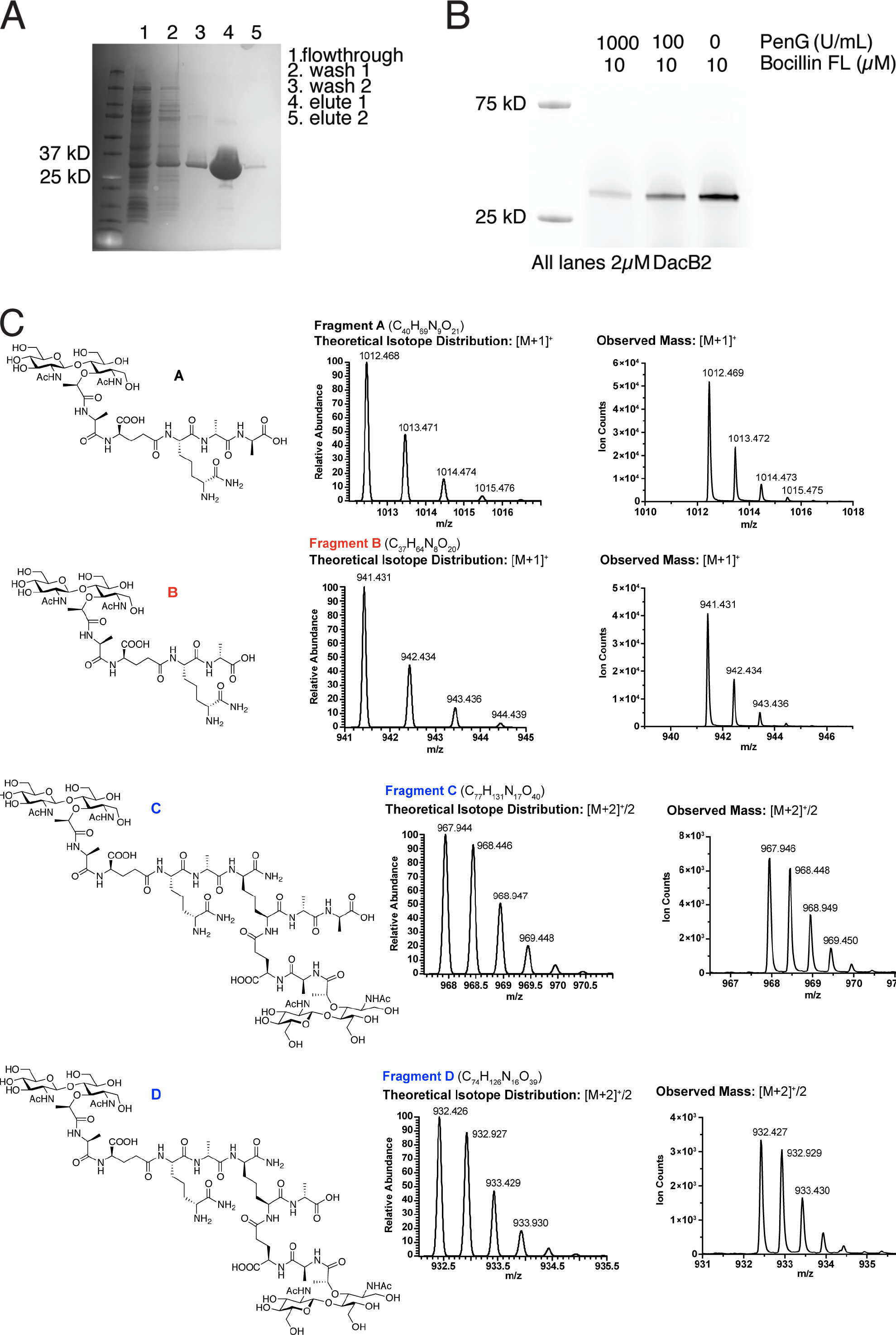
MSMEG_2433 (DacB2) functions as a D,D-carboxypeptidase and D,D-endopeptidase in vitro. **(A)** Coomassie-stained gel of purified His_6_-DacB2. **(B)** Bocillin-FL and Penicillin G binding assay of purified DacB2. **(C)** Mass spectra of the reaction products of DacB2 digestion reactions.

**Figure 5- figure supplement 1:**
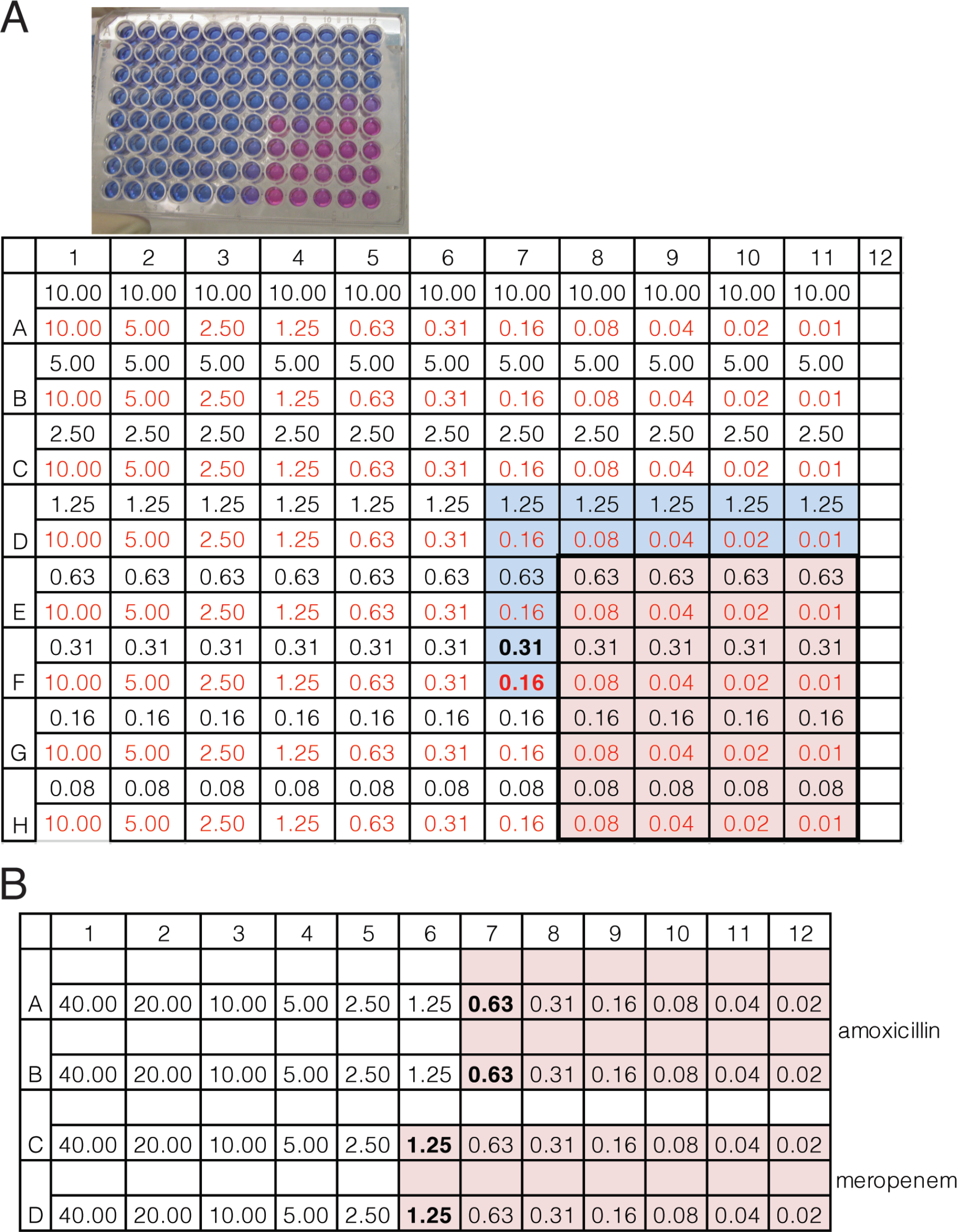
Minimum inhibitory concentration (MIC) of amoxicillin or meropenem alone or in combination against Mycobacterium tuberculosis. **(A)** Resazurin MIC plate and dilution matrix of amoxicillin and meropenem in combination in a checkerboard MIC plate. The table below shows the concentration of each drug per well in µg/mL. The concentration of meropenem is in black text and the concentration of amoxicillin is in red text in each well. **(B)** Dilution matrix of amoxicillin or meropenem (alone). The concentrations of drugs are shown in the table. Pink indicates metabolically active cells, blue indicates not metabolically active. In both single drug and checkerboard MIC plates, 5µg/mL clavulanate was used.

**Figure 5- figure supplement 2:**
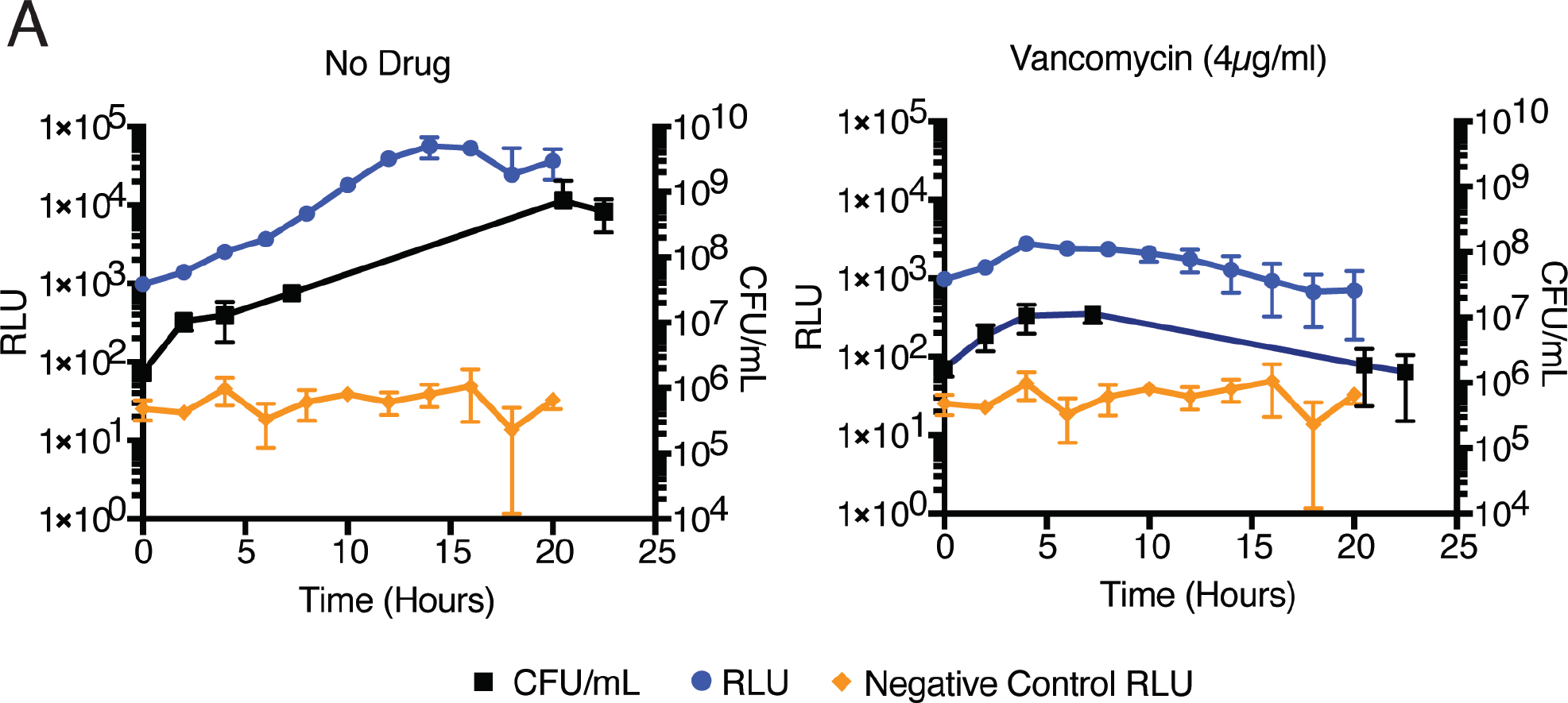
Light production (RLU) correlated to colony forming units (CFU) in mycobacterial cells expressing *luxABCDE* in drug treatment. **(A)** *Mycobacterium smegmatis* colony forming units (CFU) and luminescence (RLU) during drug treatment.

## Video captions

**Figure 2-video1- Time-lapse of ΔLDT cells in iso-osmolar media.**

This video corresponds to Figure 2A (top). This is a time-lapse microscopy video of ΔLDT Msm cells growing in 7H9 (iso-osmolar) media. Frames are 15 minutes apart, and the video is 5 frames/second.

**Figure 2-video2- Time-lapse of ΔLDT cells in high-osmolar media.**

This video corresponds to Figure 2A (bottom). This is a time-lapse microscopy video of ΔLDT Msm cells growing in 7H9 +150mM sorbitol (high-osmolar) media. Frames are 15 minutes apart, and the video is 5 frames/second.

**Figure 4-video1- Time-lapse of ponA1-RFP.**

This video corresponds to Figure 4A,B. This is a time-lapse microscopy video of Msm cells expressing ponA1-RFP growing in 7H9 media. Frames are 15 minutes apart, and the video is 5 frames/second.

**Figure 4-video2- Time-lapse of ldtE-mRFP.**

This video corresponds to Figure 4A,B. This is a time-lapse microscopy video of Msm cells expressing ldtE-mRFP growing in 7H9 media. Frames are 15 minutes apart, and the video is 5 frames/second.

**Figure 4-video3- Time-lapse of dacB2-mRFP.**

This video corresponds to Figure 4A,B. This is a time-lapse microscopy video of Msm cells expressing dacB2-mRFP growing in 7H9 media. Frames are 15 minutes apart, and the video is 5 frames/second.

**Supplementary table 1:**
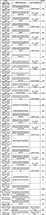
List of Primers.

**Supplementary Table 2:** List of Strains.

| Strain | Description | Primers |
| --- | --- | --- |
| KB134 | mc2155ΔldtA::loxP | 208/209; 210/211 |
| KB156 | mc2155ΔldtA::loxP + ΔldtE::zeoR | 220/221; 222/223 |
| KB200 | mc2155ΔldtA::loxP ΔldtE::zeoR + ΔldtB::hygR | 444/445; 446/447 |
| KB209 | mc2155ΔldtA::loxP ΔldtE::loxP ΔldtB::loxP + ΔldtC::hygR | 216/448; 449/219 (create original hyg KO) but used 507/508; (amplify KO from strain within the flanks) |
| KB222 | mc2155ΔldtA::loxP ΔldtE::loxP ΔldtB::loxP ΔldtC::hygR ΔldtG::zeoR | 228/454; 455/231 (create original hyg KO) but used 513/514; (amplify KO from strain within the flanks) |
| KB303 (ΔLDT) | mc2155ΔldtA::loxP ΔldtE::loxP ΔldtB::loxP ΔldtC::loxP ΔldtG::loxP ΔldtF::hygR | 224/452; 453/227 (create hyg KO) but used 511/512 (amplify KO from strain or gibson) |
| KB302 | pTetO-ldtE(MSMEG_0233)-linker-mRFP in CT94 XH (XL1-Blue) | 323A/351; 352/353 |
| KB316 (ΔLDTcomp) | [mc2155ΔldtA::loxP ΔldtE::loxP ΔldtB::loxP ΔldtC::loxP ΔldtG::loxP ΔldtF::hygR] + KB302 | |
| KB428 | *E.coli* BL21 + pet28b (dacB2) | 662/663 |

**Supplemental Table 3:** FDAA FACs screen data used for Figure 1D.

**Supplemental Table 4:** ∆LDT Tnseq data used for Figure 3A. Below are the column names with a brief description- Orf - ID of gene Name - name of gene Desc - annotation of gene Sites - number of TA sites in gene Mean Ctrl - mean insertion count averaged over TA sites and replicates for wildtype strain (mc^2^155) Mean Exp - mean insertion count averaged over TA sites and replicates for knockout strain (ΔLDT) log2FC - log-fold-change, log2(meanExp/meanCtl) Sum Ctrl - sum of insertion counts over TA sites and replicates for wildtype strain (mc^2^155) Sum Exp - sum of insertion counts over TA sites and replicates for knockout strain (ΔLDT) Delta Sum - difference of sums (sumExp-sumCtl) p-value - probability of null hypothesis (i.e. no significant difference between strains) estimated from resampling distribution Adj. p-value - p-values after applying Benjamini-Hochberg correction for multiple tests

